# Atypical developmental remodeling of dopamine neurons involves AKT-GSK3β signaling and glia activity

**DOI:** 10.1101/2025.08.29.673168

**Authors:** Xiaofan Zhang, Wei Chen, Yayu Wang, Xun Tu, Xiangyu Zhang, Ronald L. Davis, Lily Yeh Jan, Yuh Nung Jan

## Abstract

Neuronal remodeling is essential for sculpting neural circuits, and its disruption has been implicated in neurodevelopmental and neuropsychiatric disorders. Yet the molecular and cellular diversity of remodeling across neuron types remains incompletely understood. Here, we uncover a distinct remodeling mode in a subtype of *Drosophila* dopamine neurons (DANs) critical for learning, memory, sleep, and locomotion. Unlike the stereotypical pruning-then-regrowth paradigm, these DANs undergo a transient axon overgrowth followed by selective pruning during metamorphosis. Remarkably, DAN axon pruning proceeds independently of canonical ecdysone signaling and instead involves neuron-intrinsic AKT-GSK3β signaling and extrinsic glial activity. Disruption of AKT-GSK3β signaling alters microtubule stability and impairs glial recruitment and clearance of axonal debris. Notably, the role of AKT-GSK3β is cell-type specific, underscoring mechanistic diversity in remodeling programs. These findings reveal an unexpected overgrowth-then-pruning developmental trajectory, establishing DANs as a powerful model to uncover the mechanisms underlying neuronal remodeling, circuit maturation, and neurodegeneration.

## Introduction

Neuronal remodeling is a fundamental process that shapes the architecture and function of the nervous system during development and in response to experience^1–4^. Disruption of this process may contribute to a range of neurodevelopmental and neuropsychiatric disorders, including autism spectrum disorder and schizophrenia^3–7^. Growing evidence suggests that developmental neuronal remodeling and age-related neurodegeneration may share conserved molecular mechanisms, raising the possibility that similar pathways might govern neuronal degeneration across life stages^1,8,9^.

The fruit fly *Drosophila melanogaster* has long served as a powerful model for investigating the cellular and molecular basis of neuronal remodeling. A well-characterized paradigm of developmental neuronal remodeling involves the pruning of larval-stage neurites followed by regrowth to establish adult-specific circuits during *Drosophila* metamorphosis. This stereotyped pruning-then-regrowth remodeling paradigm is exemplified in many neuron types across the central nervous system (CNS) and peripheral nervous system (PNS), including cholinergic Mushroom Body (MB) neurons, GABAergic anterior paired lateral (APL) neurons, and olfactory projection neurons (PNs) in the CNS, and dendritic arborization (da) neurons in the PNS^10–14^. Whether dopamine neurons (DANs) undergo a similar remodeling trajectory was unknown.

Neuronal remodeling is driven by a combination of intrinsic and extrinsic mechanisms. The steroid hormone ecdysone functions as an essential regulator of this process across diverse neuron types that undergo pruning-then-regrowth remodeling^10–15^. Ecdysone exerts its effects through cell-autonomously expressed ecdysone receptors and downstream pathways involving caspase signaling and the ubiquitin-proteasome system (UPS)^10,13–22^. However, whether ecdysone signaling universally governs neuronal remodeling across neuron types remains largely unknown.

Pruning of neurites during neuronal remodeling typically occurs through local degeneration, followed by debris clearance mediated by extrinsic non-neuronal cells. In the CNS, glial cells engulf and eliminate degenerating neurites from MB neurons and PNs^13,23,24^. In the PNS, epidermal cells function as primary phagocytes that remove dendritic debris from da neurons^25^. Whether DANs engage in similar remodeling dynamics or rely on comparable neuron–glia interactions during development remains an open question.

Here, we identify a previously unrecognized mode of neuronal remodeling in *Drosophila* DANs. In contrast to the stereotypical pruning-then-regrowth remodeling paradigm, DANs exhibit an atypical overgrowth-then-pruning remodeling trajectory, characterized by the transient overextension of a new axon branch from an established circuit during a narrow developmental window, which is subsequently selectively pruned to establish adult-stage circuitry. Remarkably, this pruning process occurs independently of canonical ecdysone signaling, caspase activity, and the ubiquitin-proteasome system. Instead, we identify the AKT-GSK3β signaling pathway as a critical cell-intrinsic regulator of this process. We further show that DAN axon pruning proceeds through a local degeneration mechanism that involves an interplay between AKT-GSK3β signaling, microtubule cytoskeleton, and glial activities. Given the essential roles of AKT-GSK3β signaling and glial activity in DAN degeneration in Parkinson’s disease (PD), our findings raise the possibility that shared principles may link DAN pruning with disease-associated neurodegenerative processes^1,8,9,26–29^.

## Results

### Atypical developmental remodeling of dopamine neurons in *Drosophila*

To investigate whether dopamine neurons (DANs) undergo developmental remodeling, we focused on the PPL1 (protocerebral posterior lateral 1) cluster of DANs in the central brain of *Drosophila melanogaster* (fruit fly). The PPL1 cluster consists of approximately 12 DANs per hemisphere, which can be divided into five distinct subtypes with stereotyped innervation patterns across different compartments of the Mushroom Body (MB) axons^30,31^ (Figure S1A and S1B). PPL1 DANs are essential for learning and memory, forgetting, sleep, and locomotion, thus sharing several functional and mechanistic similarities with DANs in the substantia nigra pars compacta (SNc) of mammals^32–35^. Moreover, PPL1 DANs represent the only DAN subset that consistently undergoes neurodegeneration across multiple *Drosophila* Parkinson’s disease (PD) models^36–38^, making them an excellent system for investigating the mechanisms underlying DAN development in both health and disease.

We used the TH-D’-Gal4 driver to label PPL1 DANs^32^ and trace their development across various stages. While TH-D’-Gal4 labels approximately 12 DANs that project to the vertical lobe, junction, and heel regions of the MB in adult flies, we found that at the third instar larval stage, it labels only 3 DAN cell bodies, whose axons predominantly innervate the heel region (corresponding to the MP1 or PPL1γ1pedc region, hereafter referred to as MP1) (Figure S1B). This observation aligns with previous reports indicating that TH-D’-Gal4 primarily labels MP1 DANs at the third instar larval stage^39^.

During metamorphosis, the MB lobes undergo dramatic remodeling: their larval-stage-specific vertical and horizontal lobes are pruned within 18 hours after puparium formation (APF), followed by regrowth of the horizontal lobe beginning around 24 hr APF to establish adult-specific connectivity^12,19^. Notably, a pair of GABAergic APL neurons, which innervate the MB, including the MP1 region, exhibit a comparable remodeling pattern involving axon pruning and subsequent regrowth^14^. Given that both APL neurons and MP1 DANs innervate the MB, and that both APL neurons and MB itself remodel within a similar time frame^12,14,19^, we asked whether MP1 DANs follow a comparable remodeling trajectory.

Unexpectedly, we observed an overgrowth of DAN axons from the densely innervated MP1 branch into the MB’s junction region (also known as the MV1 region) between 12 hr and 18 hr APF (Figure 1A and 1D-1E). However, this newly extended branch is transient and is progressively pruned after 24 hr APF (Figure 1A and 1F-1H). Further analysis (detailed below, Figure 2) confirmed that this structure represents an overextension of axons from a single pair of MP1 DANs into the degenerating MB MV1 region (Figure 3A-3B and S3A-S3B), which we designated as the MP1o branch. Notably, the MP1o branch was not observed prior to 6 hr APF (Figure 1A-1C), suggesting that it emerges between 6 hr and 12 hr APF, and it is largely pruned by 30–36 hr APF (Figure 1A, 1G-1H).

**Figure 1.**
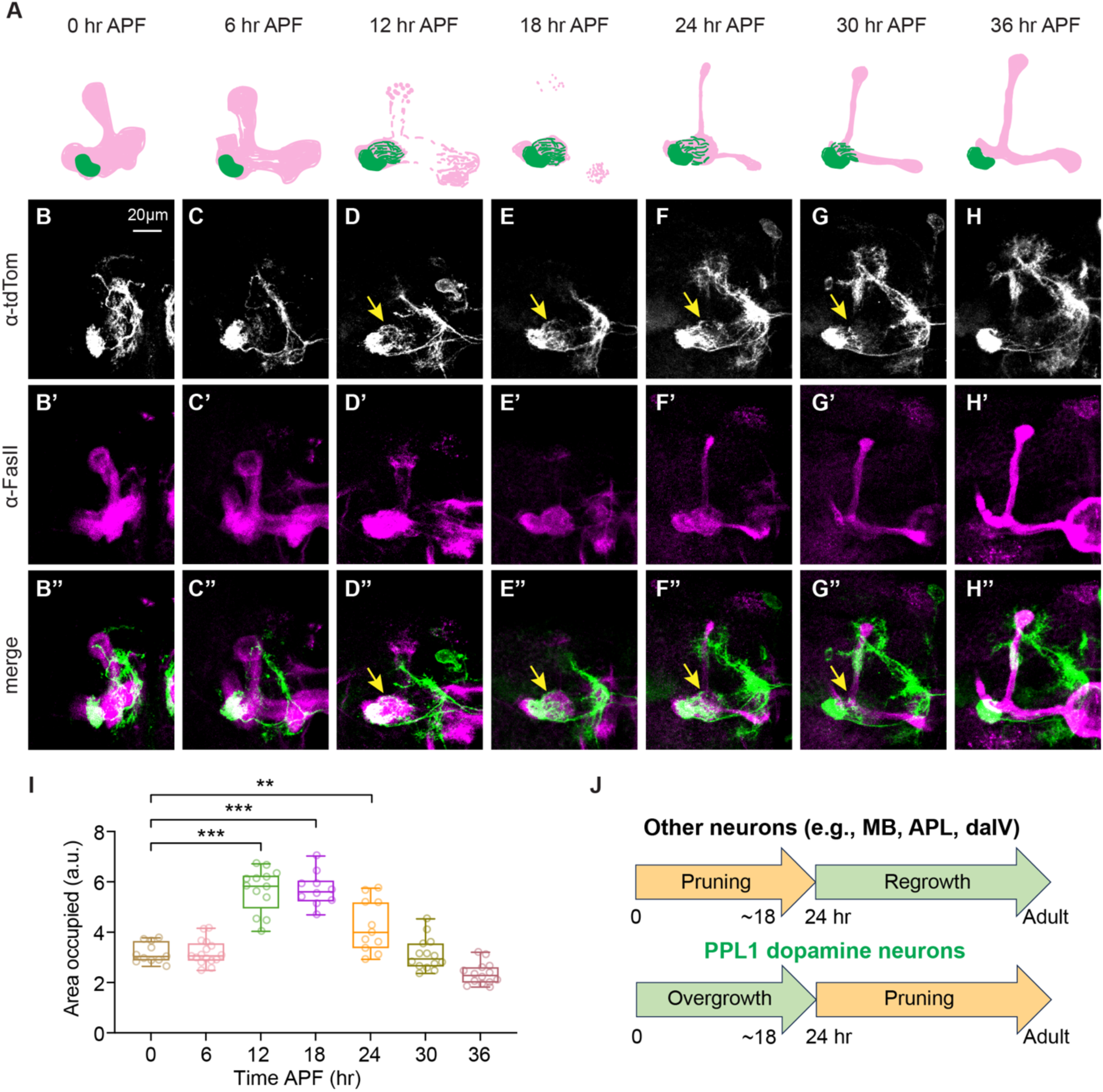
Atypical developmental remodeling of dopamine neurons. (A) Schematic of PPL1 DAN axon remodeling during metamorphosis. Green: MP1/MP1o branch; Magenta: Mushroom body (MB) lobes labeled with anti-FasII. (B-H) Confocal Z projections of brains at 0 hr (B), 6 hr (C), 12 hr (D), 18 hr (E), 24 hr (F), 30 hr (G), and 36 hr (H) APF expressing CD4-tdTomato driven by TH-D’-Gal4; MB lobes are labeled with anti-FasII (B’-H’). Yellow arrows indicate the transient MP1o branch overextending from the MP1 branch into the MB MV1 region. See also Figure S1A-S1B. (I) Quantification of the relative area occupied by MP1/MP1o branch across timepoints. \*\**p* < 0.01, \*\*\**p* < 0.001; *n* = 10-16. One-way ANOVA with Dunnett’s multiple comparisons test. (J) Summary schematic of atypical overgrowth-then-pruning developmental remodeling of DANs.

**Figure 2.**
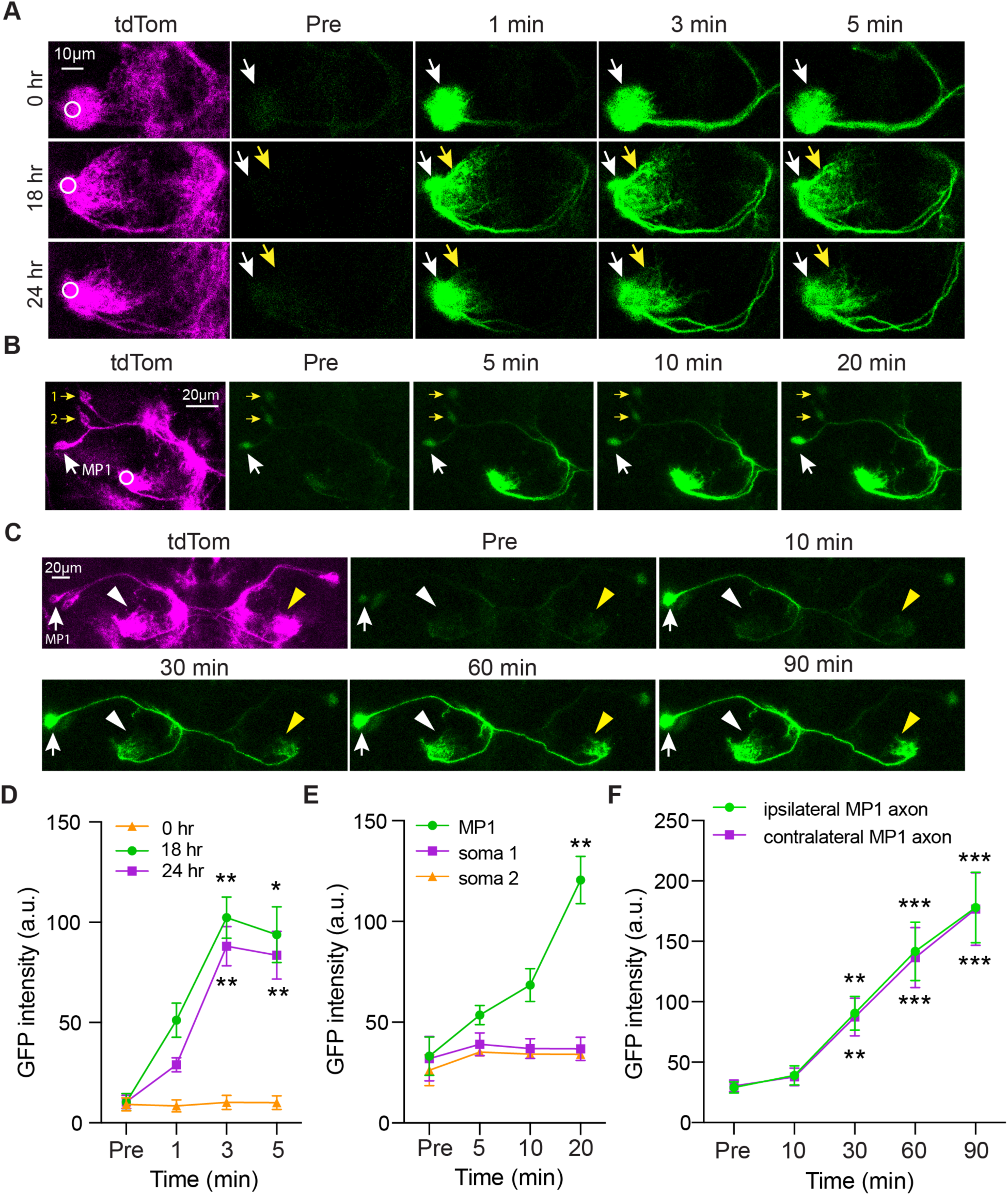
A single pair of DANs undergoes atypical developmental remodeling. (A) Photoactivation of C3PA-GFP in the MP1 branch and monitor GFP diffusion to the MP1o region at 0, 18, and 24 hr APF. White circle: ROI for photoactivation; white arrow: MP1 branch region; yellow arrow: MP1o region. (B) Photoactivation in the MP1 branch and monitor GFP diffusion to somas at 18 hr APF. White circle: ROI for photoactivation. White arrow: MP1 soma; yellow arrows: non-MP1 somas. (C) Photoactivation in MP1 soma and monitor GFP diffusion to axons at 18 hr APF. White arrow: MP1 soma; arrowheads: ipsilateral (white) and contralateral (yellow) MP1 axons. (D) Quantitation of GFP fluorescence intensity in the MP1o region in (A) before (Pre) and after (1-5 min) photoactivation. \**p* < 0.05, \*\**p* < 0.01 vs. Pre; *n* = 3-4; mean ± SEM; Repeated measures one-way ANOVA with Dunnett’s test. (E) Quantitation of GFP fluorescence intensity in MP1 and two non-MP1 somas in (B) before (Pre) and after (5-20 min) photoactivation. \*\**p* < 0.01 vs. Pre; *n* = 3; mean ± SEM; Repeated measures one-way ANOVA with Dunnett’s test. (F) Quantitation of GFP fluorescence intensity in MP1 axons in (C) before (Pre) and after (10-90 min) photoactivation. \*\**p* < 0.01, \*\*\**p* < 0.001 vs. Pre; *n* = 5; mean ± SEM; Repeated measures one-way ANOVA with Dunnett’s test.

**Figure 3.**
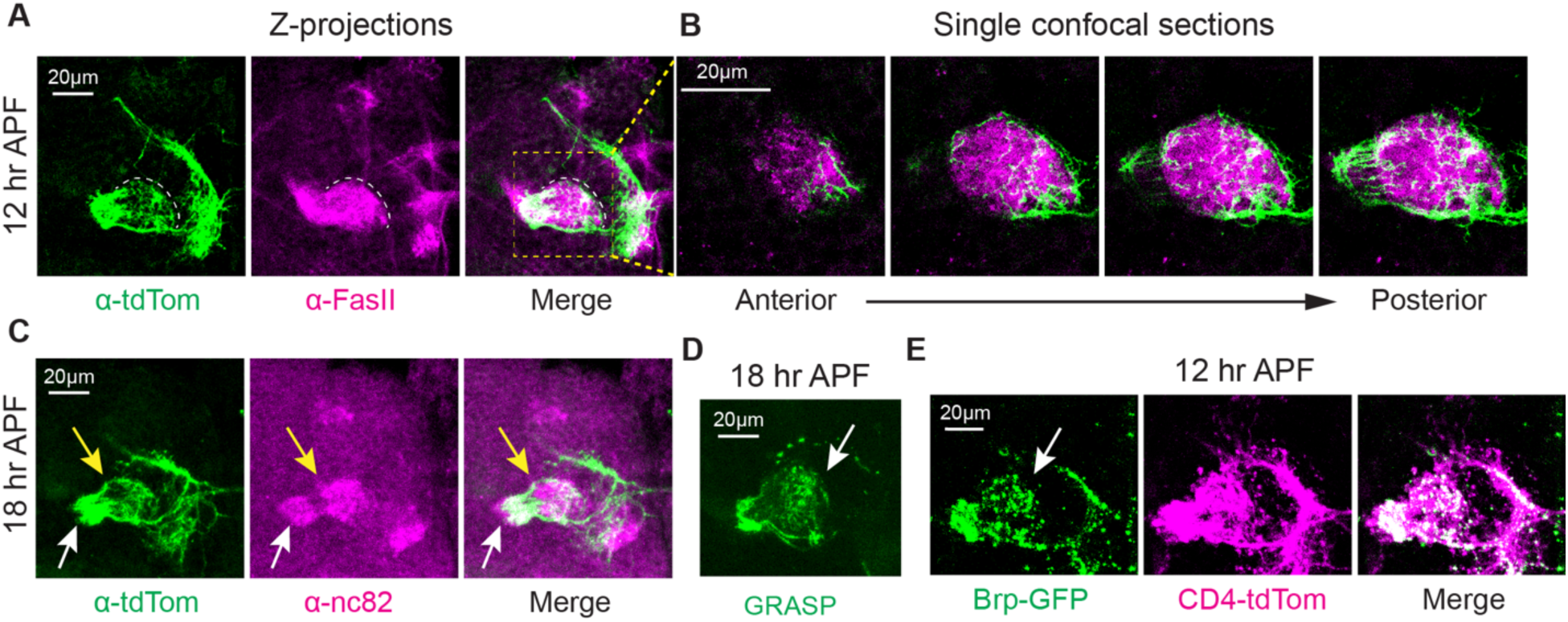
MP1o targets and is presynaptic to the MB. (A) Confocal Z projections of brains at 12 hr APF showing MP1o branch (TH-D’>CD4-tdTomato) innervating the MB. MB lobes are labeled with anti-FasII. Dashed lines mark the distal part of MP1o branch and the boundary of degenerating MB junction/MV1 region. (B) High magnification single confocal sections showing the innervation pattern of MP1o branch in MB MV1 region as in the dashed box in (A). (C) Confocal Z projections of brains at 18 hr APF expressing CD4-tdTomato driven by TH-D’-Gal4. Brain and MB lobes are visualized with anti-nc82. White arrow: MP1 region; yellow arrow: MP1o and MB MV1 regions. (D) Confocal Z projections showing GRASP signal between MP1/MP1o and MB at 18 hr APF. White arrow: MP1o region. (E) Confocal Z projections of brains at 12 hr APF expressing Brp-GFP and CD4-tdTomato driven by TH-D’-Gal4. White arrow: MP1o region.

Quantification of the relative area occupied by MP1/MP1o branches revealed a significant increase at 12, 18, and 24 hr APF compared to 0 hr APF (Figure 1I). We further validated these findings using the split-Gal4 driver MB320C, which specifically labels MP1 DANs in adult flies^40^, and found that it recapitulated the overgrowth phenotype at 18 hr APF (Figure S1C).

Together, these results demonstrate that DANs undergo developmental remodeling during metamorphosis. Whereas previous studies of developmental remodeling have documented a stereotypical pruning-then-regrowth pattern for MB neurons, PNs, APL neurons, and da neurons^10–14^, DANs exhibit an atypical overgrowth-then-pruning remodeling pattern characterized by an initial overgrowth phase followed by pruning (Figure 1J).

### A single pair of DANs undergoes atypical remodeling during metamorphosis

At both 0 and 18 hr APF, TH-D’-Gal4 labels approximately three cell bodies within the PPL1 cluster (Figure S2A-S2B, left panels). Immunostaining with anti-tyrosine hydroxylase (TH) antibodies confirmed that all three somas were TH-positive, indicating they are DANs (Figure S2A-S2B, middle and right panels). We thus asked whether the MP1 branch and the overextended MP1o branch originate from the same neuron or from distinct neurons among the three labeled DANs.

To address this, we used photoactivatable GFP (C3PA-GFP)^41^ to examine GFP diffusion between the MP1 and MP1o branches. We reasoned that if these two branches arose from the same neuron, localized photoactivation in one branch would lead to GFP diffusion into the other. Consistent with this hypothesis, photoactivation of the MP1 branch resulted in rapid GFP diffusion into the MP1o branch at both 18 and 24 hr APF (Figure 2A, 2D). In contrast, although we occasionally observed branch-like structures in the MV1 region resembling the MP1o branch at 0 hr APF, no GFP diffusion was detected between the MP1 branch and these structures (Figure 2A), confirming that the MP1o branch is not present at this earlier stage.

To further confirm that the MP1 and MP1o branches originate from a single DAN, we tracked GFP diffusion from axons in the MP1 branch back to their somas. Time-lapse imaging revealed that only one of the three DAN somas gradually accumulated GFP signal at 18 hr APF (Figure 2B, 2E), indicating it is directly connected to the MP1/MP1o branches. The remaining two DAN somas retained baseline levels of C3PA-GFP signal (Figure 2B, 2E, soma 1 and 2).

Similarly, the split-Gal4 driver MB320C labeled three somas at 18 hr APF (Figure S1C), consistent with TH-D’-Gal4 labeling. However, MB320C progressively lost labeling of the two non-MP1 DAN somas and ultimately labeled only the single MP1 DAN in the adult brain (Figure S1C, 48 hr APF and adult). The fate and function of the two non-MP1 DANs remain unknown and require further investigation. Together, these findings demonstrate that a single pair of MP1 DANs (one per brain hemisphere) undergoes atypical developmental remodeling during *Drosophila* metamorphosis.

### The MP1 DAN in each hemisphere innervates both hemispheres during development and in adulthood

Surprisingly, localized photoactivation of the MP1 branch in a single hemisphere at 18 hr APF led to GFP diffusion not only within the targeted hemisphere, but also into the soma and axons of the MP1 DAN in the contralateral hemisphere (Figure 2B, S2C). This indicates that each MP1 DAN projects its axons not only ipsilaterally within the hemisphere but also contralaterally across the midline so that both hemispheres receive its innervation during development.

To confirm this observation, we photoactivated the MP1 DAN soma in one hemisphere and observed GFP diffusion into the MP1/MP1o branches in both hemispheres (Figure 2C, 2F), further supporting the bilateral projection pattern.

We next asked whether this bilateral projection pattern persists into adulthood. Using MB320C-Gal4-driven C3PA-GFP expression in adult flies, photoactivation of a single MP1 DAN soma again resulted in GFP diffusion into MP1 branches in both hemispheres (Figure S2E-S2F), demonstrating that this bilateral connectivity is maintained beyond development. Given the established roles of MP1 DANs in memory formation and forgetting^40,42–45^, the bilateral innervation pattern may function to coordinate dopaminergic modulation across hemispheres during these cognitive processes.

### The MP1o branch includes axons presynaptic to the MB

During developmental remodeling of MB γ neurons, the larval stage γ neuron axon lobes are pruned, whereas axons in the junction (MV1) and heel/peduncle (MP1) regions are left largely intact^12,19^ (Figure 6E). The mechanisms that limit pruning to specific regions remain unknown. Subsequently, the adult-specific γ lobe regrows from the MV1/junction region^46^.

Interestingly, we found that the MP1o branch specifically targets the MV1 region of the MB, as shown by both anti-FasII and anti-nc82 (a presynaptic marker) staining at 12 and 18 hr APF (Figure 3A-3C and S3A-S3B). In fact, the MP1o branch appeared to be attracted to the MV1 region, with its distal part confined near the MV1 boundary (Figure 3A, dashed line). This innervation pattern was further confirmed in single confocal sections (Figure 3B). 3D reconstruction of the MP1/MP1o branch and MB axons reveals that the MP1o branch broadly innervates the entire MB MV1 region (Movie 1). Thus, the MP1o branch represents an overextended branch of MP1 DANs that transiently innervates the degenerating but unpruned MB MV1 region during active MB remodeling.

To test whether the MP1o branch forms synapses with the MB, we used GFP reconstitution across synaptic partners (GRASP)^47^ and observed reconstituted GFP signal in the MV1 region at 18 and 24 hr APF (Figure 3D, S3C). This finding suggests potential synaptic contacts between the MP1o branch and MB. In addition, we detected Bruchpilot:GFP (Brp-GFP), a presynaptic active zone marker^48^, in the MP1o branch at 18 and 24 hr APF (Figure 3E, S3D), indicating that the MP1o branch possesses presynaptic machinery. Together, these results suggest that the MP1o branch is likely functionally integrated with the MB.

### Local degeneration and glial involvement in MP1 DAN axon pruning

Axon degeneration is a hallmark of many neurodegenerative disorders. In PD, DAN axon degeneration often precedes soma loss, following a “dying-back” pattern in which axons degenerate progressively from distal synaptic terminals toward the cell body^49–51^. It thus appears that preventing or delaying axon degeneration could be a viable strategy to slow disease progression^1,52,53^.

Axon pruning of MB γ neurons and dendrite pruning of peripheral da neurons have been shown to occur via local degeneration^23–25,54,55^. To determine whether pruning of the MP1o branch of MP1 DANs also involves local degeneration, we examined key hallmarks of this process: axons fragmentation and phagocyte-mediated clearance of neuronal debris^1,2,19,55^.

Unlike the long axonal or dendritic tracts of MB γ neurons or class IV da (daIV) neurons^19,25^, the MP1o branch is short with densely intertwined axons (Figure 3A-3B), making it challenging to observe stereotypical fragmentation patterns. Nevertheless, we observed signs of disconnected axonal fragments at the distal end of the MP1o branch at 24-36 hr APF (Figure 1F-1H), suggesting that local degeneration may take place in the MP1o branch. In support of this notion, our data using *shi^ts^*-mediated inhibition of glial cells indicate that axon fragmentation occurs during pruning of the MP1o branch (Figure 4F, 4H).

**Figure 4.**
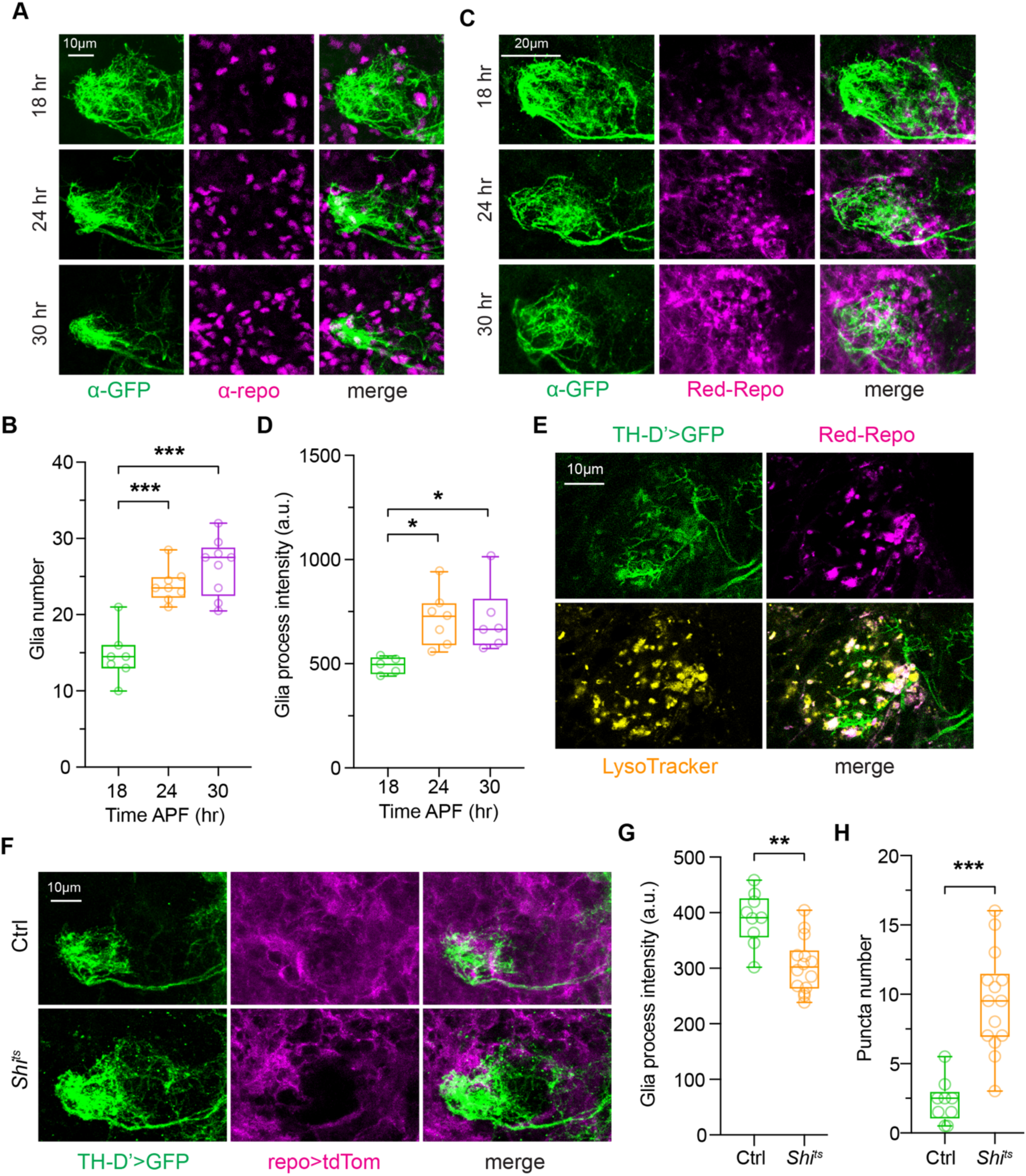
Local degeneration and glial involvement in MP1 DAN axon pruning. (A) Confocal Z projections of brains at 18 hr (top), 24 hr (middle), and 30 hr (bottom) APF expressing CD4-tdGFP driven by TH-D’-Gal4. Glial nuclei are labeled with anti-Repo. (B) Quantitation of glial cell numbers in the MP1/MP1o region in (A). \*\*\**p* < 0.001; *n* = 7-9; One-way ANOVA with Dunnett’s multiple comparisons test. (C) Confocal Z projections of brains at 18 hr (top), 24 hr (middle), and 30 hr (bottom) APF expressing mCD8-GFP driven by TH-D’-Gal4. Glial cells/processes are labeled with Red-Repo. (D) Quantitation of glial process in the MP1o region in (C). \**p* < 0.05; *n* = 5-7; One-way ANOVA with Dunnett’s multiple comparisons test. (E) Single confocal section showing MP1/MP1o branch (TH-D’>GFP), glial processes (Red-Repo), LysoTracker, and together (merge) at 24 hr APF. See also Figure S4. (F) Confocal Z projections of brains at 24 hr APF in control (top) and *shi^ts^* (bottom) groups. MP1 DANs labeled with TH-D’-GAL4>UAS-myr-GFP; glia labeled with Repo-QF2>QUAS-mtdTomato-3XHA, with or without QUAS-*shi^ts^*; Control group lacking QUAS-*shi^ts^* was subjected to the same temperature shift protocol as *shi^ts^* group. (G-H) Quantitation of glial process (G) and GFP-positive puncta (H) in MP1o region in (F). \*\**p* < 0.01, \*\*\**p* < 0.001; *n* = 9-14; Mann Whitney test.

Another hallmark of local degeneration is the involvement of phagocytes in engulfing and clearing fragmented neuronal debris. For example, glial cells have been shown to mediate MB γ neuron axon pruning by engulfing fragmented axons^23,24^. We asked whether glial cells similarly participate in pruning of the MP1o branch. To address this question, we first examined whether glial cells accumulate in the MP1/MP1o region during pruning, which would indicate their involvement. Using an anti-Repo antibody to label glial nuclei, we observed a significant increase in the number of glial cells in the MP1/MP1o branches at 24 and 30 hr APF compared to 18 hr APF (Figure 4A-4B).

Local degeneration involves active infiltration of glial processes into the degenerating neuropil^23,24,56^. To investigate this possible scenario, we used membrane-bound RFP driven by the glial-specific repo promoter (Red-Repo) to visualize glial processes in the MP1/MP1o region. In line with the observed increase in glial cell number, we detected enhanced infiltration of glial processes into the MP1/MP1o region at 24 and 30 hr APF relative to 18 hr APF (Figure 4C-4D).

Glial processes in the MP1/MP1o region exhibited punctate morphologies (Figure 4C), reminiscent of the “lumps” formed during the engulfment and degradation of degenerating γ neuron axons^24,54^. This observation raised the possibility of active glial engulfment of degenerating DAN axons, likely involving the endosomal-lysosomal pathway^23,24^. To test this possibility, we labeled endosomal-lysosomal compartments using LysoTracker and simultaneously visualized axons in the MP1/MP1o branches and glial processes. At 24 hr APF, we observed considerable number of LysoTracker-positive puncta in the MP1o region that colocalized with puncta of glial processes as well as axons in the MP1/MP1o branches (Figure 4E and S4A-S4B). These findings suggest active glial engulfment during MP1o pruning.

To further study the role of glial cells in MP1o pruning, we blocked glial endocytic activity by targeted expression of a temperature-sensitive *shibire* (*shi^ts^*) construct in glial cells (Figure 4F). *Shibire* encodes Dynamin, a GTPase essential for endocytosis, phagocytosis, and membrane trafficking^57^. *Shi^ts^*functions as a dominant-negative variant at restrictive temperatures (>29°C), thereby inhibiting endocytosis and other membrane-associated processes^58^. Previous studies have shown that *shi^ts^* expression in glia impairs glial process infiltration and delays MB γ lobe degeneration during developmental pruning^24^.

When we shifted pupae raised at the permissive temperature (23°C) to the restrictive temperature (29°C) from ∼7 hr to 24 hr APF, we observed a significant reduction in the infiltration of glial processes into the MP1/MP1o region in the *shi^ts^*group compared to control (Figure 4F-4G). Additionally, we detected an increase in axon debris, appearing as punctate structures within the MP1o region in the *shi^ts^*group relative to the control (Figure 4F and 4H), suggestive of impaired glia-mediated engulfment or degradation process. Together, these results demonstrate that pruning of MP1 DAN axons involves local degeneration and glial activity.

### Canonical ecdysone signaling is dispensable for MP1 DAN axon pruning

We next sought to investigate the cell-autonomous mechanisms underlying pruning of the MP1o branch. Ecdysone signaling is well-established as a key regulator of developmental pruning in various neuron types^10,12–15^. To test whether ecdysone signaling could similarly mediate DAN axon pruning, we expressed a dominant-negative form of the ecdysone receptor isoform B1 (EcR-B1-DN) specifically in MP1 DANs using MB320C-Gal4 and examined the fate of the MP1o branch. If ecdysone signaling were required for pruning, we would expect the MP1o branch to persist into adulthood upon EcR-B1-DN expression.

To our surprise, expression of EcR-B1-DN did not impair MP1o pruning, as axons in the MP1o branch were not detected in adult flies (Figure 5A). As a positive control validating the functionality of the construct, expression of EcR-B1-DN in MB γ neurons robustly blocked γ lobe pruning (Figure S5A). Moreover, expression of a DN form of the EcR-A isoform (EcR-A-DN) did not disrupt MP1o pruning (Figure 5B). These findings made us wonder whether MP1 DANs express significant levels of ecdysone receptors.

**Figure 5.**
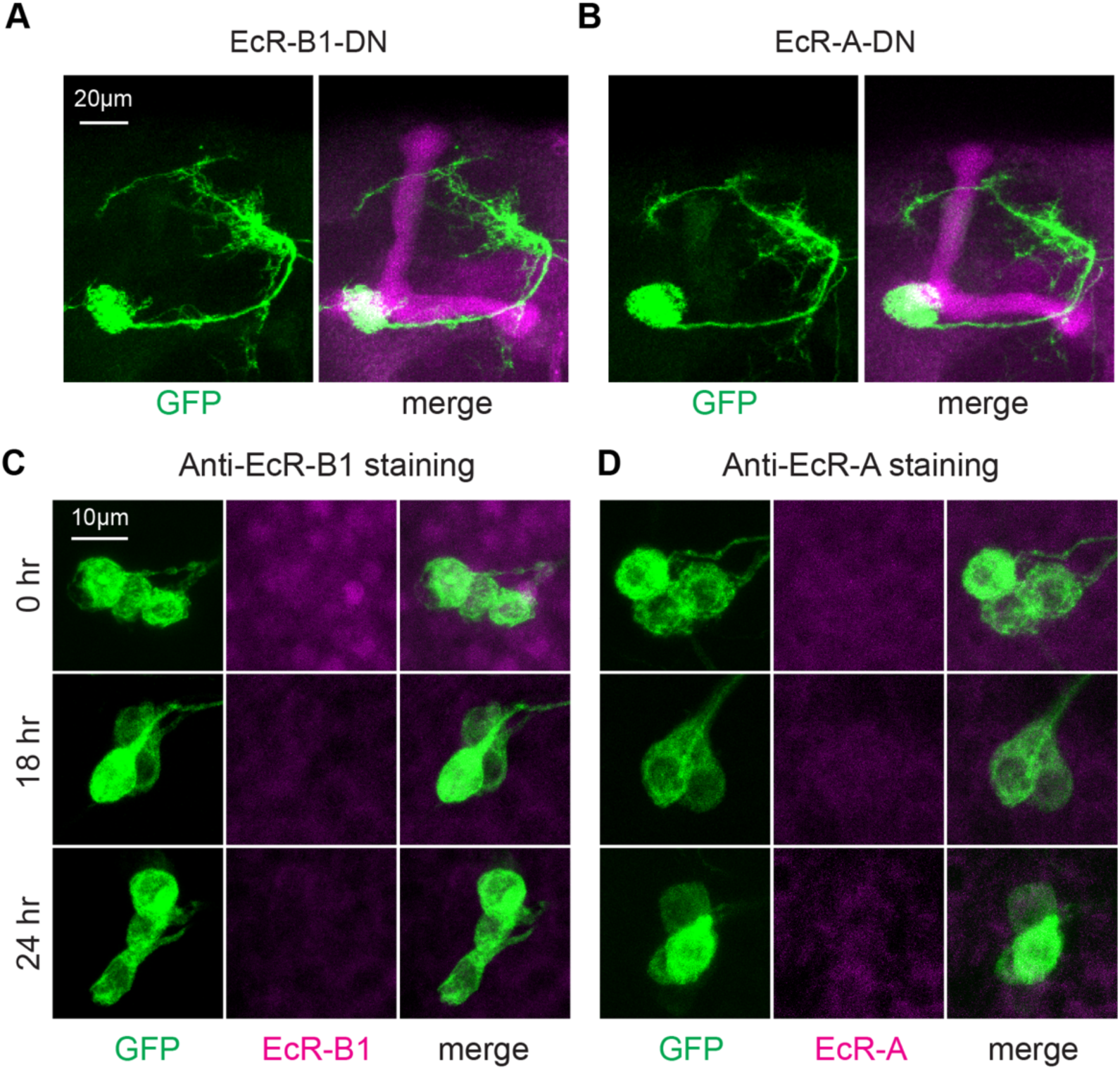
Canonical ecdysone signaling is dispensable for DAN axon pruning. (A-B) Confocal Z projections of adult brains expressing mCD8-GFP and EcR-B1-DN (A) or EcR-A-DN (B) driven by MB320C-Gal4. Magenta: MB lobes labeled with anti-FasII. No MP1o branches observed. (C-D) Confocal Z projections of brains at 0 hr (top), 18 hr (middle), and 24 hr (bottom) APF showing somas of MP1 DANs (TH-D’>mCD8-GFP) and immunostaining for EcR-B1 or EcR-A (D). EcR expression is not detected above background in MP1 DANs at any time point. See also Figure S5.

EcR-B1 or EcR-A proteins are known to be expressed in MB γ neurons, da neurons, and APL neurons at the onset of metamorphosis, where they play essential roles in developmental remodeling^10,12,14^. However, using antibody staining, we did not detect EcR-B1 or EcR-A protein expression above background levels in MP1 DAN cell bodies at 0, 18, or 24 hr APF (Figure 5C-5D). In contrast, robust EcR-B1 expression was observed in other brain regions (Figure S5B), confirming the reliability of our antibody staining. Although we cannot exclude low-level expression below detection thresholds, the absence of detectable EcR proteins in MP1 DANs is consistent with our genetic data indicating that EcR signaling is not required for MP1o pruning.

Together, these findings indicate that canonical ecdysone signaling is dispensable for pruning of the MP1o branch. While most neurons may rely on an ecdysone-dependent remodeling program, MP1 DANs appear to undergo axon pruning through an ecdysone-independent mechanism.

### Genetic screen reveals that AKT-GSK3β signaling is involved in MP1 DAN axon pruning

Although caspase activity and the UPS have been shown to be essential for developmental remodeling^17–19,21^, we found that MP1o pruning proceeds normally despite perturbations to these pathways. Specifically, expression of a dominant-negative form of the initiator caspase Dronc (Dronc-DN), as well as overexpression of the caspase inhibitors p35 or DIAP1 – manipulations known to block dendrite pruning in da neurons^17^ – did not impair MP1o pruning (Figure S5C-S5E). Similarly, RNAi knockdown of Uba1 or Cullin1 (core components of the UPS), which effectively blocks daIV dendrite pruning^21^, had no detectable effect on MP1o pruning (Figure S5F-S5G). Although we cannot exclude the possibility that the lack of pruning phenotype may result from insufficient RNAi efficiency, these findings suggest that caspase activity and UPS signaling are not essential for MP1 DAN axon pruning.

To identify molecular pathways required for DAN axon pruning, we performed a targeted genetic screen of over 200 fly lines encompassing ∼60 candidate genes implicated in neural development and remodeling. This screen revealed a critical role for GSK3β (Shaggy, Sgg in *Drosophila*) in DAN axon pruning. Expression of a dominant-negative form of GSK3β (GSK3β-DN), which mimics GSK3β inhibition, led to a pronounced pruning defect, with persistent axons in the MP1o branch detectable in adult flies (Figure 6A-6B). Given that the GSK3β activity is negatively regulated by AKT^59–61^, we asked whether elevated AKT activity similarly impacts pruning. We found that expression of a constitutively active form of AKT (AKT-CA) in MP1 DANs also disrupted MP1o pruning (Figure 6C). In contrast, expression of constitutively active S6K1, a downstream effector of mTORC1^62^, did not affect pruning (Figure 6D). These findings suggest a role for AKT-GSK3β signaling, but not mTORC1, in DAN axon pruning.

**Figure 6.**
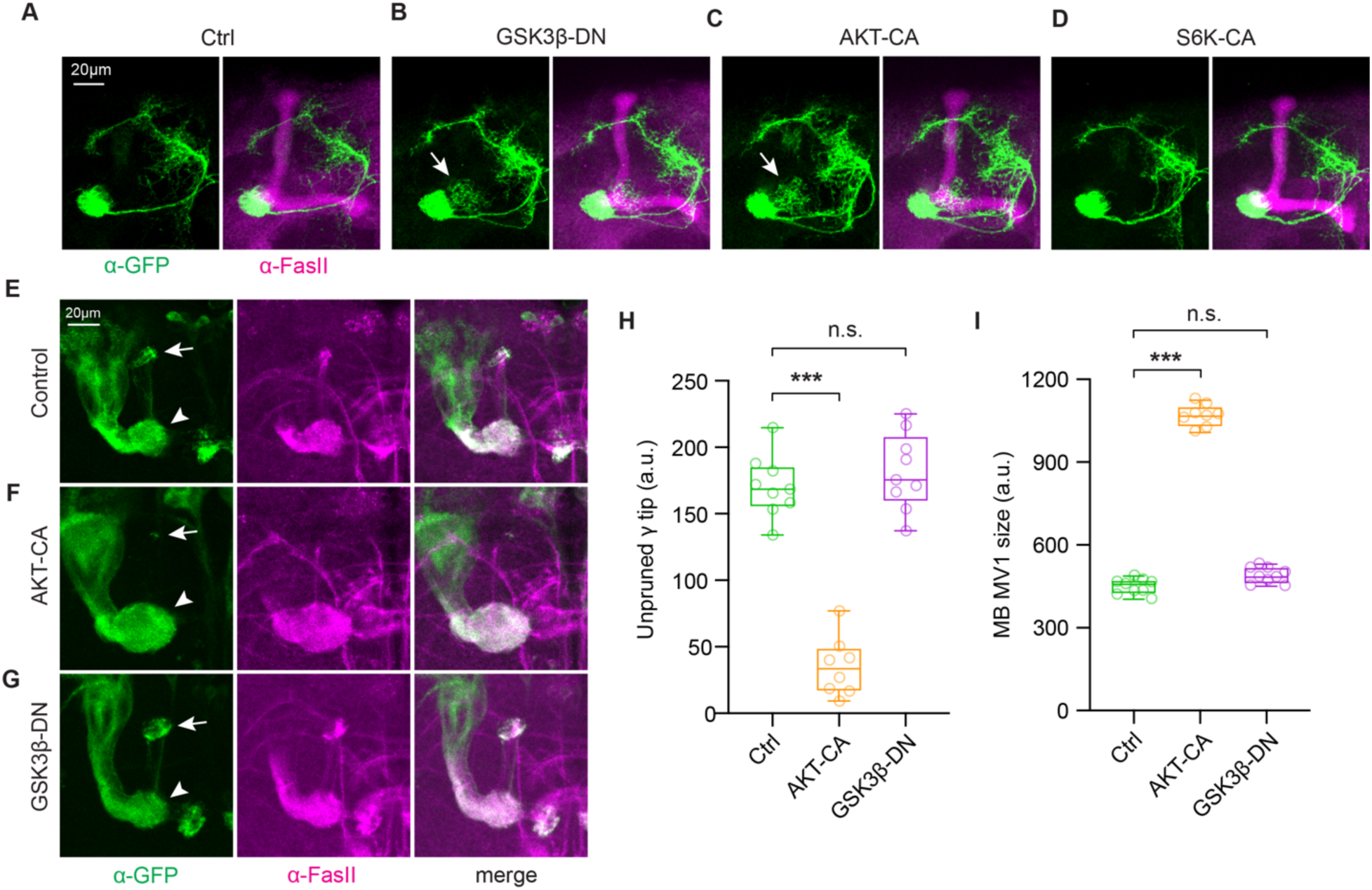
Roles of AKT-GSK3β signaling in developmental remodeling. (A-D) Confocal Z projections of adult brains expressing mCD8-GFP driven by MB320C-Gal4. Expression of GSK3β-DN (B) and AKT-CA (C), but not S6K-CA (D) in MP1 DANs impairs pruning of MP1o branch compared to control (A). MB lobes labeled with anti-FasII. White arrow indicates persistent MP1o branch in GSK3β-DN and AKT-CA groups. (E-G) Differential effects of AKT-CA (F) and GSK3β-DN (G) on MB γ neuron pruning. MB labeled with 201Y-Gal4>mCD8-GFP. White arrow: distal tip of MB γ neuron vertical lobe; white arrowhead: unpruned MB junction/MV1 region. (H-I) Quantitation of the unpruned distal tip of MB vertical lobe (H) and junction/MV1 area (I) in control (E), AKT-CA (F), and GSK3β-DN (G) groups at 18 hr APF. \*\*\**p* < 0.001; n.s., not significant; *n* = 8-9; One-way ANOVA with Dunnett’s multiple comparisons test.

### Differential roles of AKT-GSK3β signaling in neuronal remodeling

We next asked whether AKT-GSK3β signaling is similarly involved in neuronal remodeling in other neuron types, such as MB γ neurons and daIV neurons. We found that expression of AKT-CA or GSK3β-DN in MB γ neurons did not impair γ neuron axon pruning at 18 hr APF (Figure 6E-6F), consistent with previous reports that AKT is not required for γ neuron pruning^16^. Unexpectedly, AKT-CA expression appeared to enhance γ neuron axon pruning, as the distal tips of unpruned γ lobes—clearly visible in controls—was barely detectable in AKT-CA brains (Figure 6E-6F, arrow). Quantification of GFP-labeled pixels confirmed a significant reduction in signal at the distal tip of the vertical lobes in AKT-CA brains compared to wild type (Figure 6H). Additionally, we observed a significant enlargement of the MB junction/MV1 region in AKT-CA flies (Figure 6F, arrowhead, and 6I). These phenotypes were not observed in GSK3β-DN brains (Figure 6G-6I), suggesting that AKT and GSK3β may play distinct roles in γ neuron remodeling.

Although RNAi knockdown of AKT has been reported to rescue dendrite pruning defects caused by Cullin1 knockdown in daIV neurons^63^, we found that AKT-CA expression did not impair daIV dendrite pruning at 18 hr APF (Figure S6A-S6B), possibly due to context dependent differences in AKT activities. In contrast, expression of GSK3β-DN significantly disrupted daIV dendrite pruning (Figure S6A-S6B), consistent with previous findings^64^. Together, these results reveal the differential roles of AKT-GSK3β signaling in the developmental remodeling of distinct neuron types.

### Crosstalk between intrinsic AKT-GSK3β signaling and extrinsic glial activity during MP1 DAN axon degeneration

Developmental remodeling involves coordinated interactions between cell-autonomous and non-cell-autonomous mechanisms. To explore potential crosstalk between intrinsic AKT-GSK3β signaling and extrinsic glial responses during MP1o pruning, we analyzed glial dynamics across pruning stages in AKT-CA and GSK3β-DN backgrounds. At 18 hr APF, both AKT-CA and GSK3β-DN brains exhibited a reduction in the number of glial cells in the MP1/MP1o region compared to controls, while glial process infiltration remained largely unchanged (Figure S7A). By 24 hr APF, however, both glial cell number and process infiltration were significantly reduced in AKT-CA and GSK3β-DN brains (Figure 7A-7D). These reductions persisted through 30 hr APF in the GSK3β-DN group and trended toward significance in the AKT-CA group (Figure S7B). Together, these findings suggest that AKT-GSK3β signaling suppresses local glial responses during MP1 DAN axon degeneration, revealing a potential regulatory interaction between intrinsic neuronal signaling and extrinsic glial activity.

**Figure 7.**
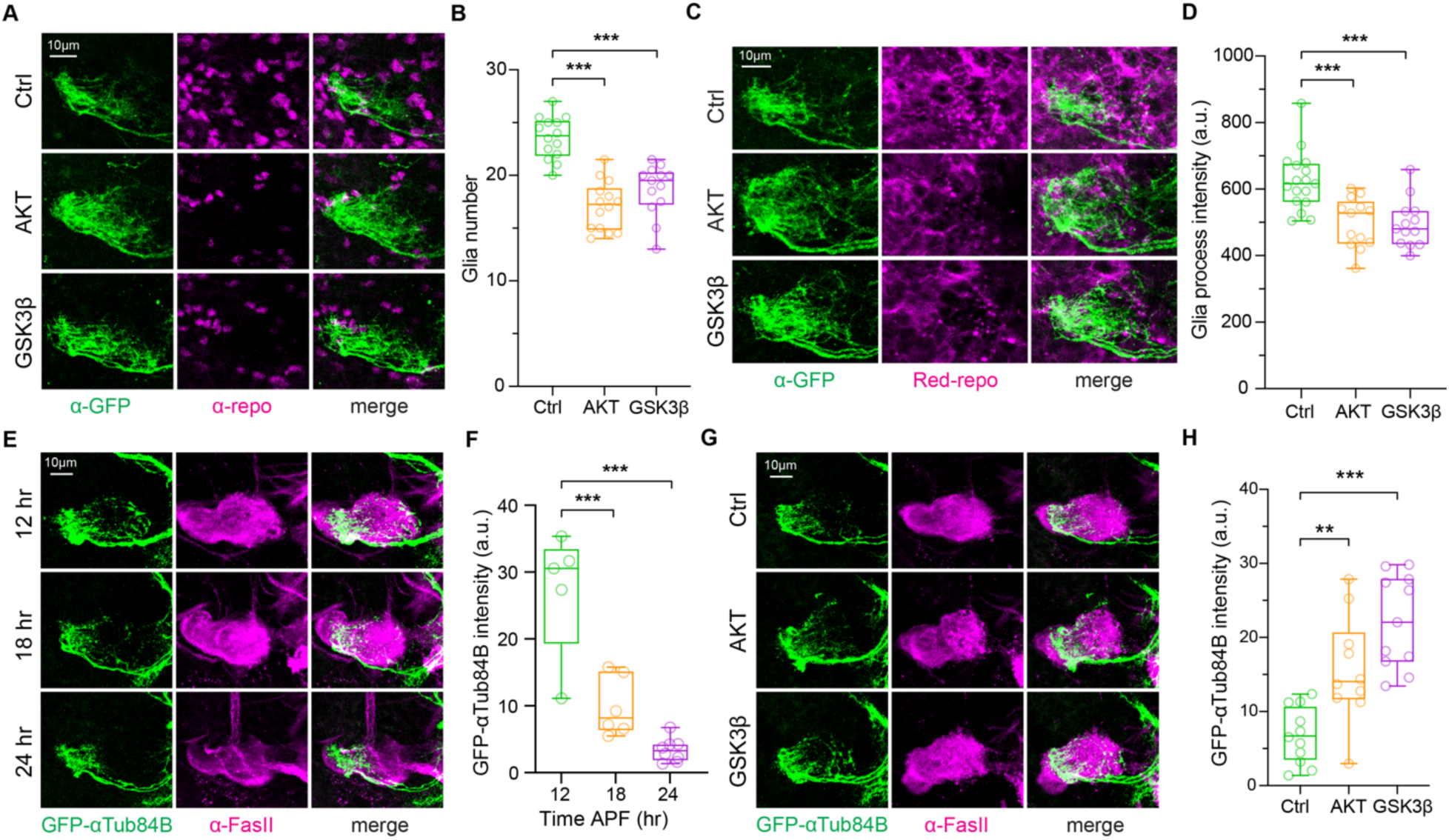
AKT-GSK3β signaling modulates microtubule cytoskeleton and glial activities during DAN axon degeneration. (A) Confocal Z projections of brains at 24 hr APF expressing mCD8-GFP driven by TH-D’-Gal4 in control (top), AKT-CA (middle), and GSK3β-DN (bottom) groups. Glial nuclei labeled with anti-Repo. (B) Quantitation of glial cell number in MP1/MP1o region in (A). \*\*\**p* < 0.001; *n* = 13-14; One-way ANOVA with Dunnett’s multiple comparisons test. (C) Confocal Z projections as in (A) but with glial cells/processes labeled with Red-Repo. (D) Quantitation of glial process in MP1o region in (C). \*\*\**p* < 0.001; *n* = 13-17; One-way ANOVA with Dunnett’s multiple comparisons test. (E) Confocal Z projections of brains at 12 hr (top), 18 hr (middle), and 24 hr (bottom) APF expressing GFP-αTub84B driven by TH-D’-Gal4. MB lobes labeled with anti-FasII. (F) Quantitation of GFP-αTub84B signal intensity in MP1o region in (E). \*\*\**p* < 0.001; *n* = 5-8; One-way ANOVA with Tukey’s multiple comparisons test. (G) Confocal Z projections of brains at 18 hr APF expressing GFP-αTub84B driven by TH-D’-Gal4 in control (top), AKT-CA (middle), and GSK3β-DN (bottom) groups. MB lobes labeled with anti-FasII. (H) Quantitation of GFP-αTub84B signal intensity in MP1o region in (G). \*\**p* < 0.01, \*\*\**p* < 0.001; *n* = 10-12; One-way ANOVA with Tukey’s multiple comparisons test.

### AKT-GSK3β signaling regulates microtubule cytoskeleton breakdown during DAN axon degeneration

To investigate how AKT-GSK3β signaling may influence DAN axon pruning, we examined its role in regulating microtubule cytoskeleton integrity, which is known to be disrupted prior to glial engulfment in other developmental pruning contexts: In both MB γ neurons and daIV neurons, early microtubule disruption is a hallmark of neural degeneration, with glial engulfment occurring only after microtubule cytoskeleton breakdown^19,54,55,64^.

We used GFP-tagged αTub84B (GFP-αTub84B) as a marker to visualize the microtubule cytoskeleton in the MP1/MP1o branches during pruning. At 12 hr APF, GFP-αTub84B signal was robustly detected in the MP1o branch, with strong expression in its distal portion (using the unpruned MB junction/MV1 region as reference) (Figure 7E). However, by 18 and 24 hr APF, the GFP-αTub84B intensity progressively diminished in the MP1o branch (Figure 7E-7F), consistent with microtubule breakdown during the early phase of axon degeneration^19,54^.

To determine whether AKT-GSK3β signaling modulates this process, we expressed AKT-CA or GSK3β-DN in MP1 DANs and quantified GFP-αTub84B levels. Both manipulations resulted in significantly elevated GFP-αTub84B signal in the distal MP1o region compared to controls (Figure 7G-7H), suggesting that AKT-GSK3β signaling stabilizes microtubule cytoskeleton during degeneration of MP1 DAN axons. Together, these findings suggest a model that AKT-GSK3β signaling cell-autonomously regulates microtubule cytoskeleton breakdown, thereby modulating glial engagement during DAN axon degeneration.

## Discussion

### Atypical developmental remodeling of DANs

This study identifies a previously uncharacterized mode of neuronal remodeling in dopamine neurons (DANs) of *Drosophila melanogaster*. In contrast to the stereotypical pruning-then-regrowth remodeling paradigm observed in most neuron types – including MB neurons, APL neurons, PNs, and daIV neurons^10–14^ – MP1 DANs follow an atypical overgrowth-then-pruning trajectory (Figure 1 and S8). This remodeling trajectory is particularly striking given that both MP1 DANs and APL neurons innervate the MB, yet exhibit opposite temporal patterns of remodeling. This atypical mode of remodeling of DAN axons represents a notable deviation from established paradigms and underscores the heterogeneity and complexity of neuronal remodeling programs in the developing brain.

The MP1o branch emerges within a narrow developmental window and targets the MB MV1, a region known to be spared during MB γ neuron pruning^12,19^. The MP1o branch exhibits presynaptic markers and potential synaptic contacts with MB, suggesting that it may be functionally integrated, albeit transiently. These observations raise intriguing possibilities: MP1 DAN axons in the MP1o branch may transiently modulate MB circuit activity, provide scaffolding for subsequent MB connectivity, or serve as a checkpoint to ensure MB pruning or regrowth fidelity.

### Pruning independent of canonical ecdysone signaling

Our results indicate that MP1 DANs prune independently of the canonical ecdysone– EcR signaling pathway (Figure 5). This is in stark contrast to most remodeling neurons in *Drosophila*, including MB neurons, PNs, APL neurons, da neurons, and thoracic ventral (Tv) neurons, which show EcR expression and rely on ecdysone signaling to initiate remodeling^10,12–15^. A recent study reports that pruning of Beat-Va_M_ neurons in the ventral nerve cord (VNC) of *Drosophila* is also ecdysone-independent, although EcR expression is detected in these neurons^65^. In contrast, our data indicate that neither EcR-B1 nor EcR-A is detectable in MP1 DANs. While we cannot fully exclude the possibility that MP1 DANs may respond to ecdysone at levels below our detection threshold or with distinct sensitivity compared to other neuron types, our results nonetheless argue that pruning of MP1 DANs proceeds through mechanisms distinct from the canonical ecdysone-EcR pathway. Together, these findings point to neuron-type–specific pruning triggers and underscore the diversity of remodeling logic across the nervous system^1,2,8,16^.

### Role of AKT-GSK3β signaling in DAN axon pruning

Our genetic screen revealed a critical role for AKT-GSK3β signaling in DAN remodeling (Figure 6). Inhibition of GSK3β or constitutive activation of AKT impairs pruning of the MP1o branch, supporting a model in which the proper regulation of AKT and GSK3β activity is essential for timely DAN axon elimination.

Mechanistically, we find that AKT-GSK3β signaling modulates microtubule cytoskeleton breakdown during pruning. Pruning of the MP1o branch is accompanied by progressive microtubule disassembly, a hallmark of early axonal degeneration. Expression of AKT-CA or GSK3β-DN preserves microtubule integrity and inhibits pruning, indicating that cytoskeletal destabilization is a prerequisite for MP1 DAN axon degeneration. Our findings support a model in which AKT-GSK3β regulates the pruning process by modulating microtubule destabilization—a known prerequisite for local axon degeneration^2,19,54^.

### Context-dependent roles of AKT-GSK3β in neuronal remodeling

The role of AKT-GSK3β signaling in neuronal remodeling appears to be neuron type-specific. This pathway is involved in MP1 DAN axon pruning but is dispensable in MB γ neurons and only partially required in da neuron remodeling (Figure 6). These findings highlight the context-dependent deployment of shared signaling pathways across distinct neuronal populations. AKT-GSK3β signaling may thus serve specialized functions depending on the developmental timing, neurite architecture, and extrinsic cellular environment of the remodeling neurons^2,8,16,20,55,66^.

### Involvement of glial cells in DAN axon pruning

In parallel to cell-intrinsic mechanisms, we demonstrate that pruning of MP1 DAN axons involves local degeneration and glial activity (Figure 4). Glial cells accumulate in the MP1/MP1o region during the pruning window and extend processes into the region with degenerating axons. Lysosomal markers are enriched within glial processes and colocalize with DAN axon fragments, suggestive of active phagocytosis of degenerating neurites. Inhibition of glial endocytic function via expression of temperature-sensitive *shi^ts^* impairs pruning and leads to accumulation of axonal debris, consistent with a requirement for glial engulfment in axon clearance. These observations are similar to those in MB γ neuron and da neuron pruning, where glial or epidermal cells act as phagocytes for debris engulfment and clearance^23–25,54^.

### Crosstalk between intrinsic and extrinsic remodeling mechanisms

We find that AKT-GSK3β signaling modulates not only axon-intrinsic cytoskeletal remodeling but also glial behavior (Figure 7). Expression of AKT-CA or GSK3β-DN in MP1 DANs suppresses both glial cell accumulation and glial process infiltration into the MP1o region. This suggests that cytoskeletal integrity within axons may act as a permissive cue for glial engagement, either by modulating the presentation of phagocytic cues on the axonal membrane or by influencing neuron-glia signaling dynamics. This is consistent with observations in MB γ neurons and da neurons, where microtubule breakdown precedes glial engulfment and may serve as a prerequisite for efficient clearance^23,54,55^. Future studies will be needed to dissect whether glia actively sense microtubule disassembly or respond to other neuron-derived “find-me” or “eat-me” signals that are modulated by AKT-GSK3β signaling^67–69^.

### Implications for neurodegenerative disease

The identification of AKT-GSK3β and glia-involved local axon degeneration mechanism in DAN remodeling has important implications for neurodegenerative diseases. In PD, DAN axon degeneration often precedes soma loss, following a “dying-back” pattern^49,70–72^. Activation of AKT or inhibition of GSK3β has been shown to suppress axon degeneration and soma loss of DANs in mouse models of PD^26–29^. In addition, AKT promotes axon regeneration through inhibition of GSK3β or by modulating microtubule dynamics^73–75^. Growing evidence also underscores the critical role of glial cells in the pathogenesis of neurodegenerative disorders, including Alzheimer’s disease (AD), PD, and Huntington’s disease (HD)^76–78^.

Together, these findings suggest that developmental pruning mechanisms may be co-opted or dysregulated in pathological contexts. Although not the primary focus of this study, our results reveal that DANs are capable of extending new axonal branches even after circuit formation (i.e., during the MP1o overgrowth phase). Elucidating the molecular and cellular mechanisms underlying this axon overgrowth may inform novel therapeutic strategies aimed at promoting DAN axon regeneration in PD.

In conclusion, this study defines a novel mode of neuronal remodeling in *Drosophila* DANs, integrating AKT-GSK3β–regulated microtubule destabilization with glial engulfment to drive local DAN axon degeneration. These findings broaden the mechanistic repertoire of developmental pruning and provides a framework for investigating neuron-glia interactions in both healthy development and disease-related neurodegeneration.

## Acknowledgments

We thank Y.C Chang, Y. Guo, K. Li, N. C. Noyes, C.F. Teo, A. Yong, X. Ming, M. Zubia, C. Chen, P. Jin, J. Ko, and F. Xie for critical discussion; members of the Jan lab for support and feedback throughout this study; We thank S. Younger, X. Zhai, T. Cheng, and M. Y. L. Fontaine for assistance with fly work and reagents; We also thank C. Han, Janelia Research Campus, and the Bloomington *Drosophila* Stock Center for providing fly lines. This work was supported by NIH grants to Y.N.J. (R35NS097227 and R35NS137312), L.Y.J. (R35NS122110), and R.L.D. (5R35NS097224). Y.N.J. and L.Y.J. are Howard Hughes Medical Institute investigators.

## Author contributions

Y.N.J. and L.Y.J. supervised the project. X.Z. conceived and initiated the project while in the laboratory of R.L.D., and continued designed and performed experiments in the laboratories of Y.N.J. and L.Y.J. X.Z. performed immunostainings, the genetic screen, photoactivation experiments, and data analysis. W.C. assisted with experiments in da neurons. Y.W. contributed to sample preparation. X.Y.Z. assisted with the genetic screen. X.T. contributed to data analysis. X.Z. wrote the manuscript with input from all authors.

## RESOURCE AVAILABILITY

Further information and requests for resources and reagents should be directed to the lead contact Yuh Nung Jan.

## Materials availability

All unique reagents generated in this study are available from the lead contact.

## Data and code availability

This study does not generate datasets or code.

## EXPERIMENTAL MODEL AND STUDY PARTICIPANT DETAILS

### *Drosophila* husbandry and strains

Flies were maintained on standard *Drosophila* food at 25°C, 70% relative humidity, under a 12/12 hr light/dark cycle unless otherwise specified. Pupae were collected at 2 hr intervals and aged to defined developmental stages for subsequent experiments. The following fly strains were used: w1118, TH-D’-Gal4 driver^32^, MB320C-Split-Gal4 driver^40^, UAS-CD4-tdTomato^79^, UAS-CD4-tdGFP^79^, UAS-mCD8-GFP^80^, UAS-myr-GFP (BDSC #32197), UAS-myrGFP.QUAS-mtdTomato-3xHA (BDSC #77479), UAS-C3PA-GFP^41^, MB-spGFP11 construct^81^, UAS-spGFP1-10 construct^47^, UAS-Brp-GFP (gift from Stephan Sigrist), RedRepo^82^, Repo-QF2 (BDSC #66477), QUAS-shi^ts1^ (BDSC #30013), UAS-EcR-B1^DN^ (BDSC# 6872), UAS-EcR-A^DN^ (BDSC #9452), UAS-AKT^CA^ (BDSC #80935), UAS-GSK3β^DN^ (BDSC #5360), UAS-S6K^CA^ (BDSC #6913), 201y-Gal4 (BDSC #64296), PPK-Gal4 driver^79^, UAS-GFP-αTub84B (BDSC #7373), UAS-Dronc^DN^ (BDSC #58992), UAS-p35 (BDSC #5073), UAS-DIAP1 (BDSC #6657), TRiP RNAi control lines (BDSC# 36303 or 36304), UAS-Uba1-RNAi (BDSC #76066), UAS-Cullin1-RNAi (BDSC #29520). Genotypes of flies in each experiment are described in Table S1.

### Immunostaining

Drosophila brains were dissected and processed as previously described^42,83^. Briefly, brains were dissected in cold S2 medium (Life Technologies, Cat# 21720-024) and fixed overnight at 4°C in 1% paraformaldehyde in S2 medium. Following fixation, brains were washed with PAT3 buffer [phosphate buffered saline (PBS) with 0.5% BSA and 0.5% TritonX-100], blocked in 3% normal goat serum (NGS) in PAT3, and incubated with primary antibodies diluted in blocking buffer (3% NGS in PAT3) for 3 hours at 23°C, followed by overnight incubation at 4°C. After washing with PAT3, brains were incubated with secondary antibodies in blocking buffer for 3 hours at 23°C and then at 4°C for an additional 5 days. Brains were then washed with PAT3 and subsequently with 1XPBS before mounting with mounting medium (Vector Laboratories, Cat# H-1000-10). Primary antibodies used were: rabbit anti-GFP (1:1000, Invitrogen, Cat# A11122), chicken anti-GFP (1:1000, Abcam, Cat# ab13970), rabbit anti-Dsred (1:1000, Takara Bio, Cat# 632496), rabbit anti-TH (1:500, Millipore, Cat# AB152), mouse monoclonal anti-FasII (1:100, DSHB, 1D4, RRID:AB_528235), mouse monoclonal anti-nc82 (1:50, DSHB, RRID:AB_2314866), mouse monoclonal anti-repo (1:100, DSHB, 8D12, RRID:AB_528448), mouse monoclonal anti-EcR-B1 (1:50, DSHB, AD4.4, RRID:AB_2154902), mouse monoclonal anti-EcR-A (1:50, DSHB, 15G1a, RRID:AB_528214). Secondary antibodies used were: Alexa 488 goat anti-rabbit (1:800, Invitrogen, Cat# A11008), Alexa 488 goat anti-chicken (1:800, Invitrogen, Cat# A11039), Alexa 594 donkey anti-mouse (1:500, Jackson Labs, Cat# 715-585-150), Alexa 633 goat anti-rabbit (1:800, Invitrogen, Cat# A21070), Alexa 633 goat anti-mouse (1:800, Invitrogen, Cat# A21052). Images were collected on a Leica SP8 confocal microscope using either a 10X dry or 40X oil objective, at 1024 X 1024 or 2048 X 2048-pixel resolution, with excitation lasers at 488, 594, and/or 633 nm. Image analysis was performed using Fiji/ImageJ.

### LysoTracker staining

Drosophila brains were dissected in S2 medium at room temperature and transferred to 100 nM LysoTracker Deep Red (Invitrogen, Cat# L12492) diluted in S2 medium. Samples were incubated for 15 min at room temperature, then washed three times with PBS before imaging.

### Photoactivation of C3PA-GFP

To label MP1 or MP1o branch with C3PA-GFP, brains were dissected in 1XPBS and transferred to fresh 1XPBS on a microscope slide with a donut-shaped spacer. A coverslip was then applied to the spacer to minimize PBS evaporation and prevent brain movement during subsequent photoactivation and imaging. Prior to photoactivation, MP1 and/or MP1o branches were imaged using a 10X dry objective at 1024 X 1024-pixel resolution with 488 nm (for C3PA-GFP) and/or 561 nm (for tdTomato) laser excitation (designated as the “Pre” image). Photoactivation was performed on a Leica SP8 microscope using a 405 nm laser. TdTomato signal was used to identify the MP1 or MP1o region, and a circular ROI of the same size across experiments was drawn within the boundaries of the MP1 or MP1o region for photoactivation. C3PA-GFP was photoactivated by scanning the ROI with 4 iterations, each lasting 1.471seconds. In experiments requiring visualization of GFP diffusion from axons to soma or soma to axons, the photoactivation protocol was repeated every 10 or 20 minutes to enhance GFP signal for diffusion. After photoactivation, the MP1, MP1o, and/or soma were imaged at defined time points using the same imaging settings as that for the “Pre” image.

### *Shi^ts^* experiments

To inhibit dynamin function using *shi^ts^*, pupae cultured at 23°C were shifted to 29°C from approximately 7 hr APF to 24 hr APF. Pupae were dissected at 24 hr APF immediately after removal from 29°C, then fixed and stained as described above.

### 3D rendering

Three-dimensional reconstruction of the MP1/MP1o branch and MB axons (Video S1) was performed using Imaris software (version 10.1, Bitplane).

### Quantification and statistical analysis

Confocal images were analyzed using Fiji/ImageJ, and statistical analyses were performed in GraphPad Prism 9. The sample size for each group, *p*-values and the statistical tests employed for each experiment are indicated in the figures or figure legends.

To quantify the relative area occupied by MP1/MP1o (Figure 1), Z-projections were generated from confocal image stacks, and brightness and contrast were adjusted uniformly across all developmental stages. Regions within the MP1/MP1o area with signal above a defined threshold were outlined and measured in Fiji/ImageJ. The resulting area was then normalized to brain size, estimated as the distance between the left and right MP1 branches, to calculate the relative area occupied by MP1/MP1o.

To quantify C3PA-GFP fluorescence intensity before and after photoactivation (Figure 2, S2), a circular ROI of the same size was drawn around the outer boundaries of the MP1, MP1o, or soma region for each corresponding area, and the mean intensity was used for quantification.

To quantify the number of glial cells in the MP1/MP1o region, Z-projections were generated from confocal stacks. A circular ROI of the same size across all developmental stages was drawn around the outer boundaries of the MP1 and MP1o region, and the number of glial cells within the ROI was counted using Fiji/ImageJ.

To quantify glial process intensity, a Z-projection consisting of 15 optical slices (15 μm total) spanning the anterior-posterior extent of the MP1o branch was generated from confocal stacks. A circular ROI of the same size was drawn within the MP1o boundaries, and the mean intensity within the ROI was measured using Fiji/ImageJ.

To quantify GFP puncta in the *shi^ts^* experiments (Figure 4F and 4H), a Z-projection spanning the anterior-posterior extent of the MP1o branch was generated from confocal stacks and converted to greyscale in Fiji/ImageJ. Two duplicate images were created for each projection and subjected to Gaussian blur filter with Sigma values of 1 and 2, respectively. The image with Sigma = 2 was then subtracted from the image with Sigma = 1 using the Image Calculator tool. Thresholding was then applied using the MaxEntropy algorithm, which we found to be the most reliable for distinguishing true GFP puncta from background and false-positive signals. Only GFP puncta located in the distal region of the MP1o branch were quantified, as puncta in the proximal region were predominantly false positives.

To quantify the size of unpruned γ lobe vertical tips and MB peduncles (Figure 6), Z-projections were generated from confocal stacks, and brightness and contrast were adjusted uniformly across all groups. Regions corresponding to the γ lobe vertical tip or MB peduncle with signal above a defined threshold were outlined and measured using Fiji/ImageJ.

To quantify dendrite length (Figure S6), the Simple Neurite Tracer (SNT) plugin in Fiji/ImageJ was used. The total length of all traced dendrites per brain was quantified.

To quantify GFP–αTub84B intensity (Figure 7), a Z-projection consisting of 15 optical slices (15 μm total) spanning the anterior-posterior extent of the MP1o branch was generated from confocal stacks. A rectangular ROI of the same size across all experimental groups was placed around the distal part of the MP1o branch using the unpruned MB peduncle as reference. The mean intensity within the ROI was measured using Fiji/ImageJ after background subtraction using an identical ROI placed in a background region of the image.

---

**Table S1.**

| **Figure** | **Panel** | **Genotype** |
| --- | --- | --- |
| Figure 1 | B-I | TH-D'-GAL4/+; UAS-CD4-tdTomato/+ |
| Figure 2 | A-F | TH-D'-GAL4/UAS-C3PA-GFP; UAS-CD4-tdTomato/+ |
| Figure 3 | A-C | TH-D'-GAL4/+; UAS-CD4-tdTomato/+ |
|  | D | TH-D'-GAL4/+; MB-spGFP11, UAS-spGFP1-10/+ |
|  | E | TH-D'-GAL4/+; UAS-Brp-GFP/UAS-CD4-tdTomato |
| Figure 4 | A, B | TH-D'-GAL4/+; UAS-CD4-tdGFP/+ |
|  | C, D | UAS-mCD8-GFP/+; TH-D'-GAL4/Red-Repo |
|  | E | TH-D'-GAL4/Red-Repo; UAS-CD4-tdGFP/+ |
|  | F-H | UAS-myr-GFP, QUAS-mtdTomato-3xHA/+; TH-D'-GAL4/+ or QUAS-shi<sup>ts</sup>; Repo-QF2/+ |
| Figure 5 | A | UAS-mCD8-GFP/+; UAS-EcR-B1-DN/+; MB320C-GAL4/+ |
|  | B | UAS-mCD8-GFP/+; UAS-EcR-A-DN/+; MB320C-GAL4/+ |
|  | C, D | UAS-mCD8-GFP/+; TH-D'-GAL4/+ |
| Figure 6 | A | UAS-mCD8-GFP/+;; MB320C-GAL4/+ |
|  | B | UAS-mCD8-GFP/+;; MB320C-GAL4/UAS-AKT-CA |
| | C | UAS-mCD8-GFP/+;; MB320C-GAL4/UAS-GSK3β-DN |
|  | D | UAS-mCD8-GFP/+;; MB320C-GAL4/UAS-S6K-CA |
| | E-I | 201Y-GAL4, UAS-mCD8-GFP/+, UAS-AKT-CA, or UAS-GSK3β-DN |
| Figure 7 | A-D | UAS-mCD8-GFP/+; TH-D'-GAL4/Red-Repo; +/+, UAS-AKT-CA, or UAS-GSK3β-DN |
| | E, F | TH-D'-GAL4/+; UAS-GFP-αTub84B/+ |
| | G, H | TH-D'-GAL4/+; UAS-GFP-αTub84B/+, UAS-AKT-CA, or UAS-GSK3β-DN |
| Figure S1 | B | TH-D'-GAL4/+; UAS-CD4-tdTomato/+ |
|  | C | MB320C-GAL4/UAS-CD4-tdTomato |
| Figure S2 | A, B | UAS-mCD8-GFP/+; TH-D'-GAL4/+ |
|  | C, D | TH-D'-GAL4/UAS-C3PA-GFP; UAS-CD4-tdTomato/+ |
|  | E, F | UAS-C3PA-GFP/+; MB320C-GAL4/UAS-CD4-tdTomato |
| Figure S3 | A, B | TH-D'-GAL4/+; UAS-CD4-tdTomato/+ |
|  | C | TH-D'-GAL4/+; MB-spGFP11, UAS-spGFP1-10/+ |
|  | D | TH-D'-GAL4/+; UAS-Brp-GFP/UAS-CD4-tdTomato |
| Figure S4 | A, B | TH-D'-GAL4/Red-Repo; UAS-CD4-tdGFP/+ |
| Figure S5 | A | 201Y-GAL4, UAS-mCD8-GFP/UAS-EcR-B1-DN |
|  | B | UAS-mCD8-GFP/+; TH-D'-GAL4/+ |
|  | C | UAS-mCD8-GFP/+; UAS-Dronc-DN/+; MB320C-GAL4/+ |
|  | D | UAS-mCD8-GFP/+;; MB320C-GAL4/UAS-p35 |
|  | E | UAS-mCD8-GFP/+;; MB320C-GAL4/UAS-DIAP1 |
|  | F | UAS-mCD8-GFP/+;; MB320C-GAL4/UAS-Uba1-RNAi |
|  | G | UAS-mCD8-GFP/+;; MB320C-GAL4/UAS-Cullin1-RNAi |
| Figure S6 | A, B | PPK-GAL4, UAS-CD4-tdTomato/+, UAS-AKT-CA, or UAS-GSK3β-DN |
| Figure S7 | A, B | UAS-mCD8-GFP/+; TH-D'-GAL4/Red-Repo; +/+, UAS-AKT-CA, or UAS-GSK3β-DN |

