## Supplemental figures for "Atypical developmental remodeling of dopamine neurons involves AKT-GSK3β signaling and glia activity"

### Supplementary Figures

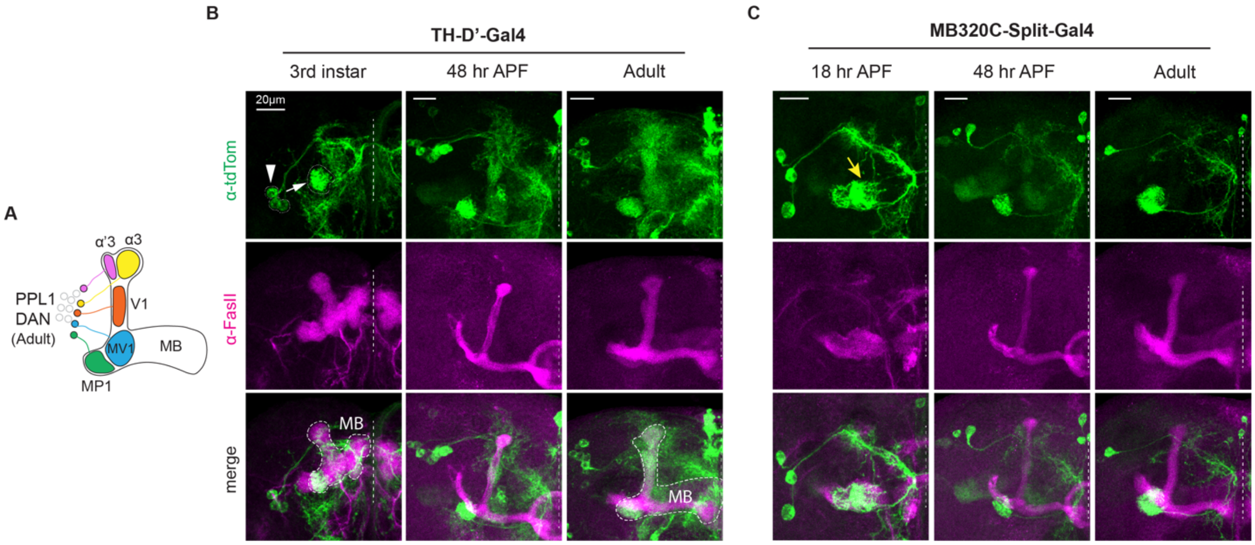


**Figure S1. Atypical developmental remodeling of DANs, related to Figure 1**

(A) Schematic of PPL1 DAN innervation patterns in the adult brain.

(B) Confocal Z projections of brains from 3rd instar larvae (left), 48 hr APF (middle), and adults (right) expressing CD4-tdTomato driven by TH-D'-Gal4. MB lobes are labeled with anti-FasII. White arrow: MP1 branch; white arrowhead: somas; dashed circles: MB.

(C) Confocal Z projections of brains at 18 hr (left), 48 hr (middle), and adult (right) expressing CD4-tdTomato driven by MB320C-Split-Gal4. Yellow arrow: MP1o branch at 18 hr APF.

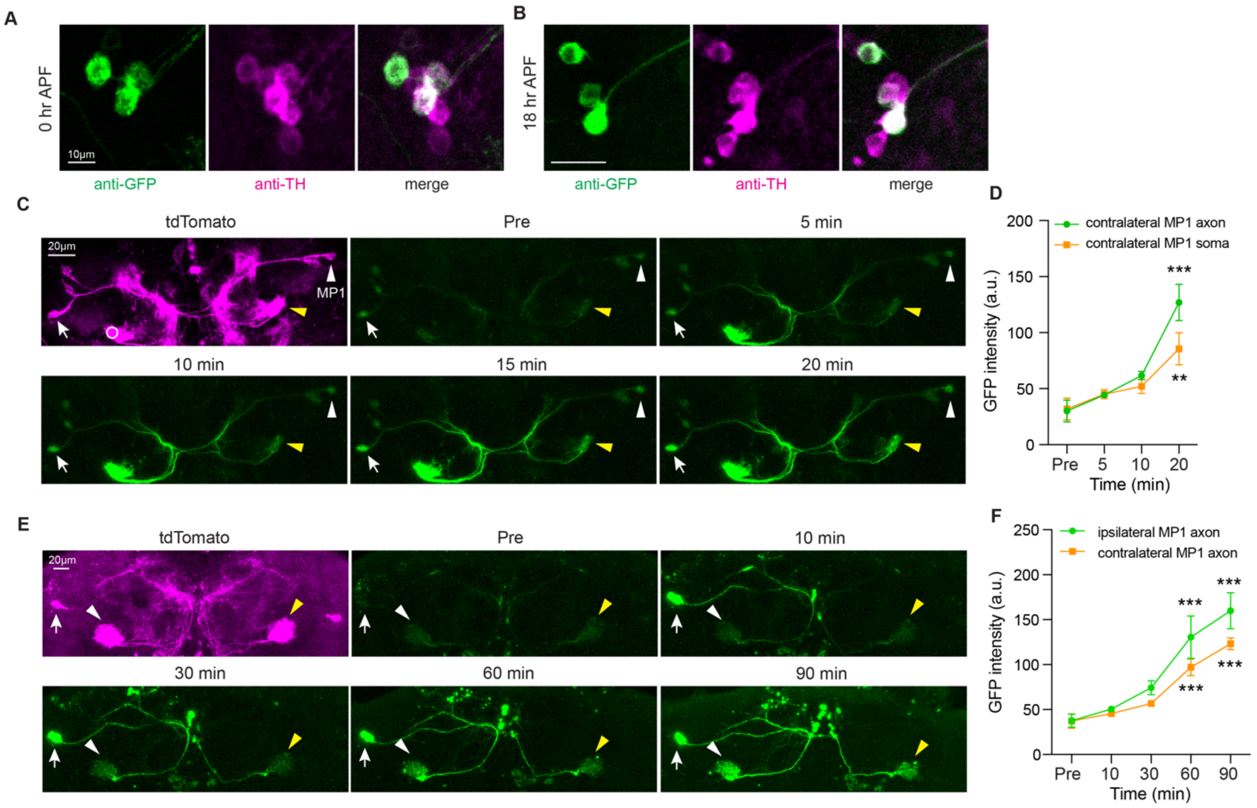


**Figure S2. A single pair of DANs undergoes atypical developmental remodeling, related to Figure 2**

(A-B) Anti-TH immunostaining confirms that all three TH-D'-Gal4-labeled somas are TH-positive at 0 hr (A) and 18 hr (B) APF.

(C) Photoactivation of C3PA-GFP in MP1 branch and monitor GFP diffusion to contralateral MP1 axon and soma (same brain as Figure 2C). White circle: ROI for photoactivation. White arrow: ipsilateral MP1 soma; white arrowhead: contralateral MP1 soma; yellow arrowhead: contralateral MP1 axon.

(D) Quantitation of GFP fluorescence intensity in contralateral MP1 axon and soma in (C) before (Pre) and after (5-20 min) photoactivation. $^{**}p < 0.01$, $^{***}p < 0.001$ vs. Pre; $n = 3$; mean ± SEM; Repeated measures one-way ANOVA with Dunnett's test.

(E) Photoactivation in MP1 soma and monitor GFP diffusion to ipsilateral and contralateral MP1 axons in adult. White arrow: ipsilateral MP1 soma; white arrowhead: ipsilateral MP1 axon; yellow arrowhead: contralateral MP1 axon.

(F) Quantitation of GFP fluorescence intensity in ipsilateral and contralateral MP1 axons in (E) before (Pre) and after (10-90 min) photoactivation. ***$p < 0.001$ vs. Pre; $n = 4$; mean ± SEM; Repeated measures one-way ANOVA with Dunnett’s test.

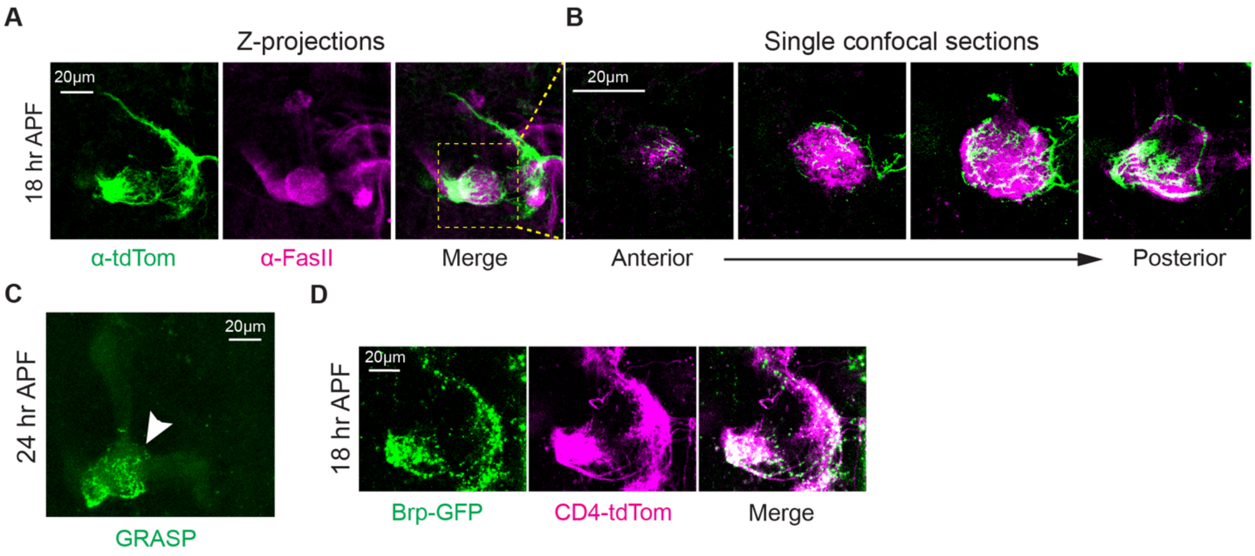


**Figure S3. MP1o targets and is presynaptic to the MB, related to Figure 3**

(A) Confocal Z projections of brains at 18 hr APF expressing CD4-tdTomato driven by TH-D'-Gal4 with MB lobes labeled by anti-FasII.

(B) High magnification single confocal sections showing the innervation pattern of MP1o branch in MB MV1 region as in the dashed box in (A).

(C) Confocal Z projections showing GRASP signal between MP1/MP1o and MB at 24 hr APF. White arrowhead: MP1o region.

(D) Confocal Z projections of brains at 18 hr APF expressing Brp-GFP and CD4-tdTomato driven by TH-D'-Gal4.

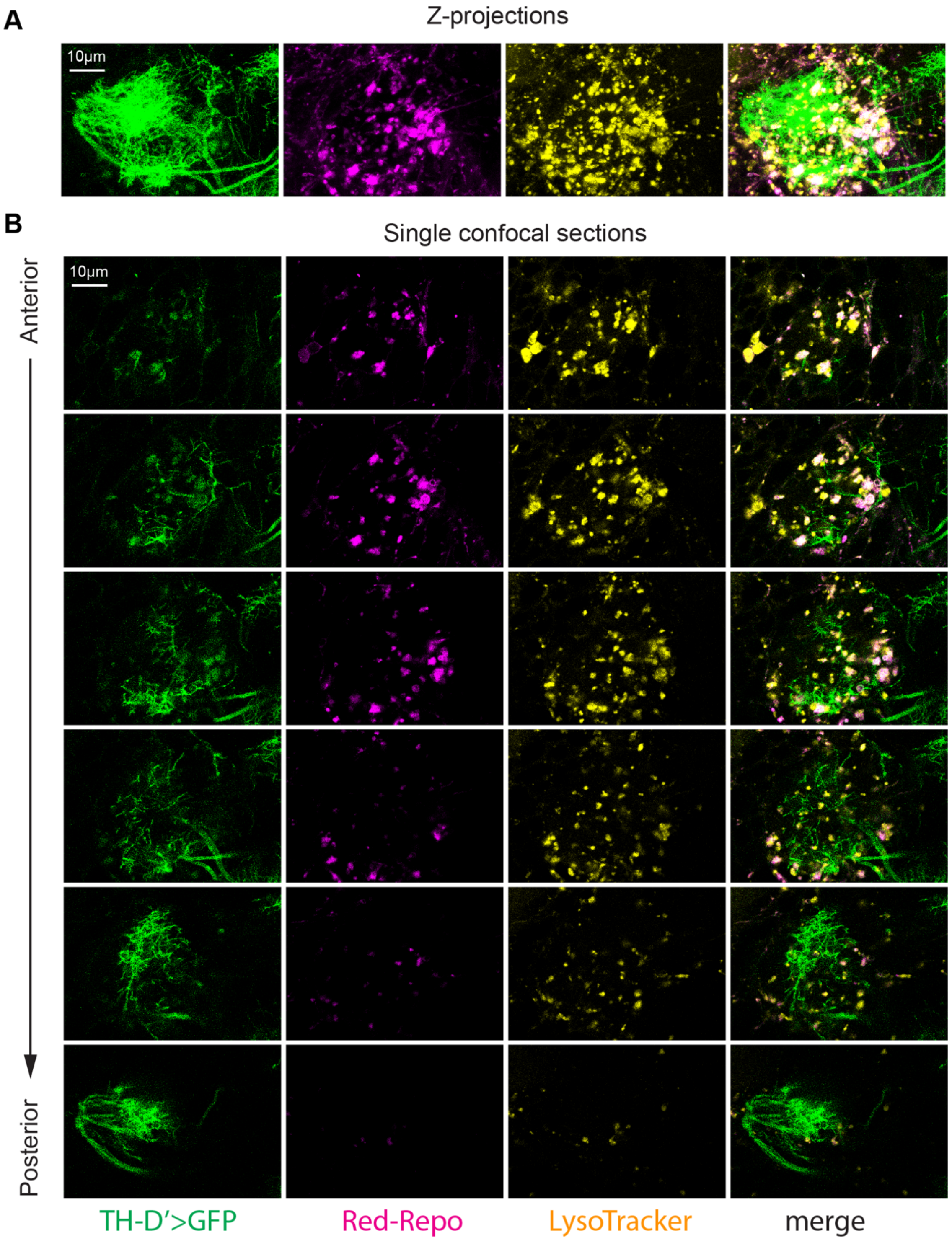


**Figure S4. Glial involvement in MP1 DAN axon pruning, related to Figure 4**

(A-B) Confocal Z projections (A) and single confocal sections (B) showing MP1/MP1o branch (TH-D’>GFP), glial processes (Red-Repo), LysoTracker, and together (merge) at 24 hr APF.

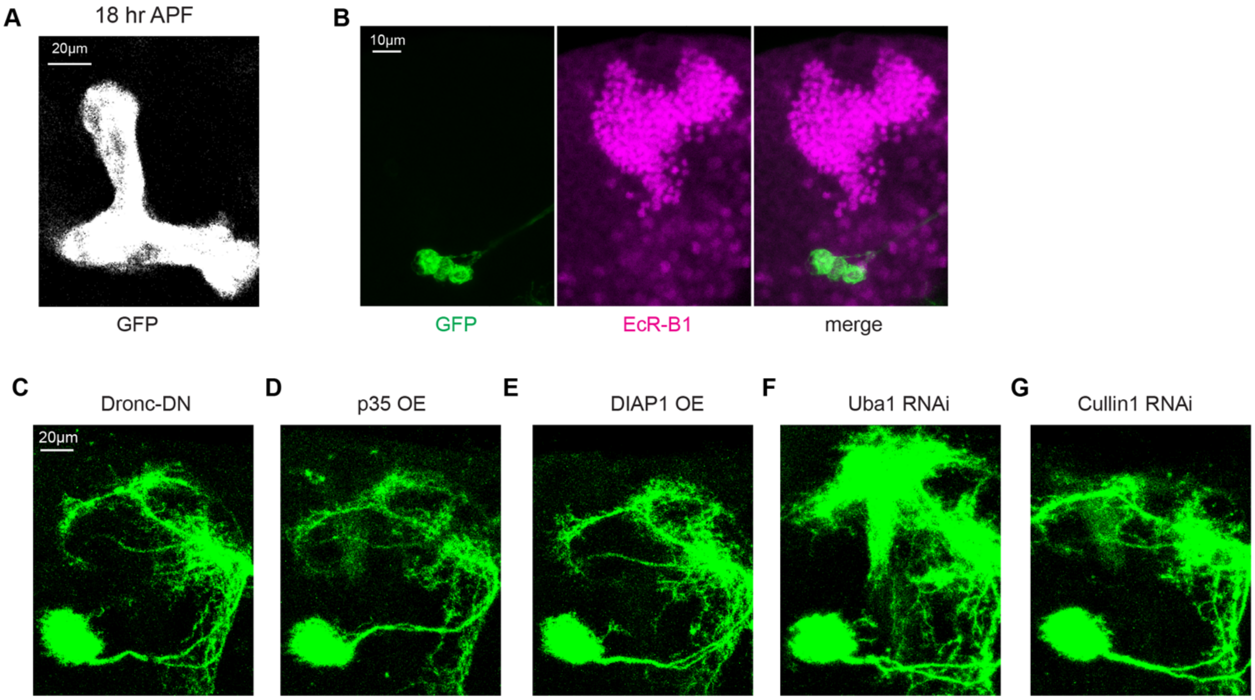


**Figure S5. Canonical ecdysone signaling, caspase activity, and UPS are dispensable for DAN axon pruning, related to Figure 5**

(A) Confocal Z projections of MB lobes at 18 hr APF expressing mCD8-GFP and EcR-B1-DN driven by 201Y-Gal4.

(B) EcR-B1 protein expression in TH-D'>mCD8-GFP brains at 0 hr APF.

(C-G) Expression of Dronc-DN (C), overexpression of p35 (D) and DIAP1 (E), and RNAi knockdown of Uba1 (F) and Cullin1 (G) do not impair MP1o pruning. Adult stage MP1 DANs are visualized with MB320C-Gal4 driving mCD8-GFP.

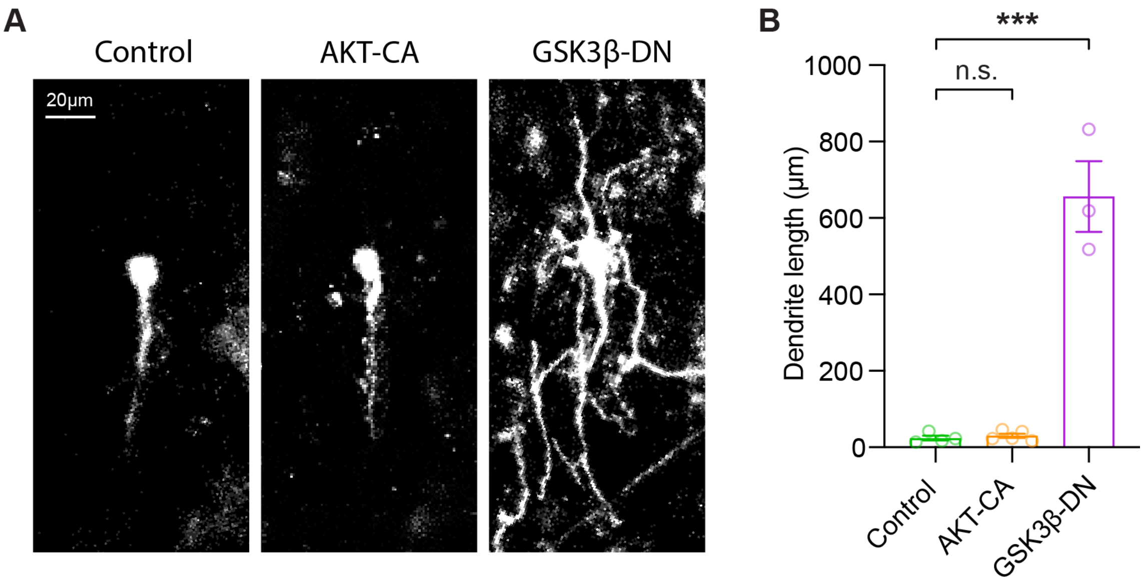


**Figure S6. Differential roles of AKT and GSK3β in dendrite pruning of daIV neurons, related to Figure 6**

(A) Confocal Z projections of daIV neurons expressing CD4-tdTomato driven by PPK-Gal4 at 18 hr APF in control (left), AKT-CA (middle), and GSK3β-DN (right) groups.

(B) Quantitation of total dendrite length in (A). ***$p < 0.001$; n.s., not significant; $n = 3\text{-}5$; mean ± SEM; One-way ANOVA with Dunnett's multiple comparisons test.

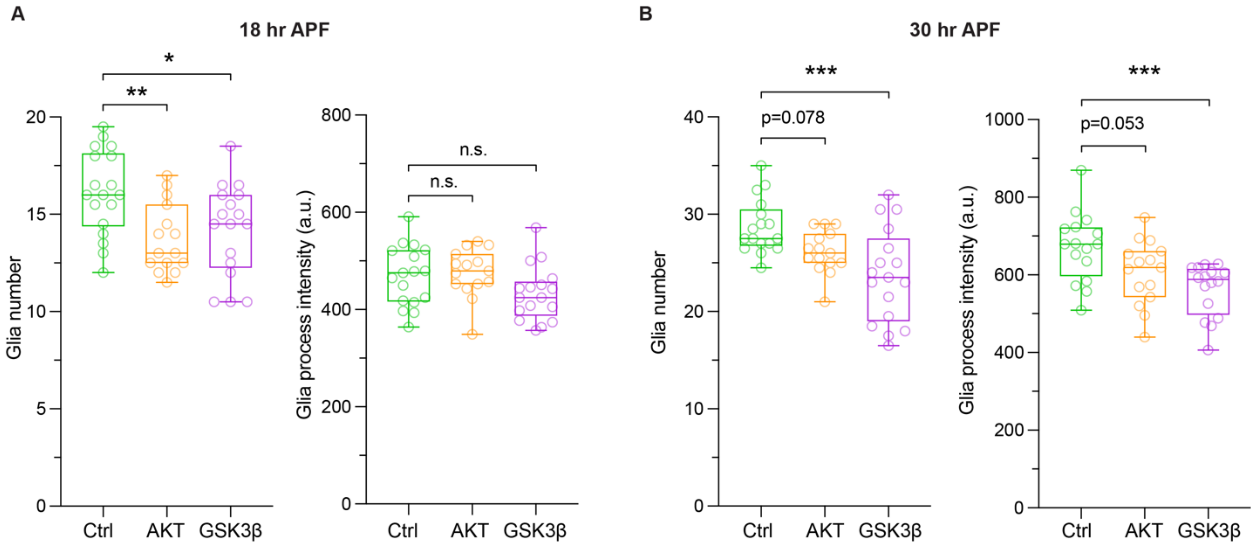


**Figure S7. Quantification of glial dynamics during DAN axon degeneration, related to Figure 7**

(A) Quantitation of glial cell number (left) and process intensity (right) in the MP1/MP1o region in control, AKT-CA, and GSK3β-DN groups at 18 hr APF. *$p < 0.05$, **$p < 0.01$; $n = 15\text{-}18$; One-way ANOVA with Dunnett's multiple comparisons test.

(B) Quantitation of glial cell number (left) and process intensity (right) in the MP1/MP1o region in control, AKT-CA, and GSK3β-DN groups at 30 hr APF. ***$p < 0.001$; $n = 15\text{-}17$; One-way ANOVA with Dunnett's multiple comparisons test.

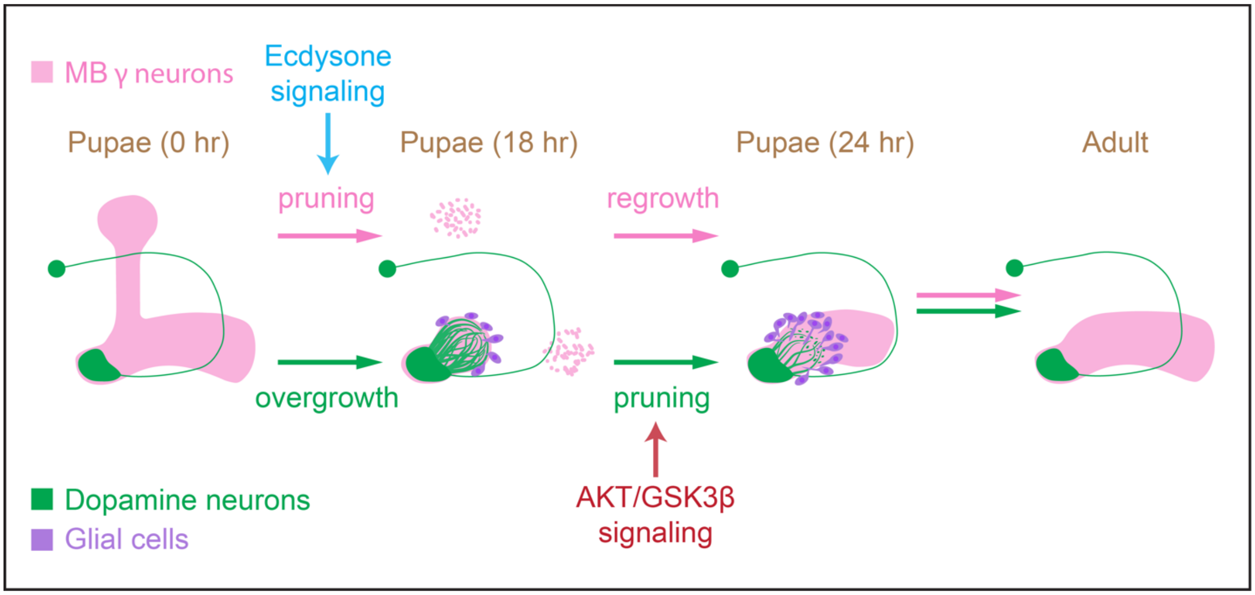


**Figure S8. Model**

In contrast to the stereotypical pruning-then-regrowth remodeling paradigm observed in most neuron types – such as MB γ neurons (illustrated here), APL neurons, PNs, and daIV neurons – MP1 DANs, which innervate the MB heel region, undergo an atypical overgrowth-then-pruning trajectory during *Drosophila* metamorphosis. Moreover, pruning of MP1 DAN axons occurs independently of the canonical ecdysone-EcR signaling pathway and instead involves neuron-intrinsic AKT-GSK3β signaling and extrinsic glial activity.

**Video S1. 3D reconstruction of MP1/MP1o branch and MB axons at 12 hr APF, related to Figure 3A.**

3D rendering of MP1/MP1o branch and MB lobes at 12 hr APF. Green: MP1/MP1o axon branches (TH-D’>CD4-tdTomato); magenta: MB peduncle (MP1) and junction/MV1 region (anti-FasII). MP1o projects into MB MV1 region.
